# Overexpression screen of chromosome 21 genes reveals modulators of Sonic hedgehog signaling relevant to Down syndrome

**DOI:** 10.1101/2022.05.19.492735

**Authors:** Anna J. Moyer, Fabian-Xosé Fernandez, Yicong Li, Donna K. Klinedinst, Liliana D. Florea, Yasuhiro Kazuki, Mitsuo Oshimura, Roger H. Reeves

## Abstract

Dysregulation of Sonic hedgehog (SHH) signaling may contribute to multiple Down syndrome-associated phenotypes, including cerebellar hypoplasia, congenital heart defects, craniofacial and skeletal dysmorphologies, and Hirschsprung disease. Granule cell precursors isolated from the developing cerebellum of Ts65Dn mice are less responsive to the mitogenic effects of SHH than euploid cells, and a single postnatal dose of the SHH pathway agonist SAG rescues cerebellar morphology and performance on learning and memory tasks in Ts65Dn mice. SAG treatment also normalizes expression levels of *OLIG2* in neural progenitor cells derived from human trisomy 21 iPSCs. However, despite evidence that activating SHH signaling can ameliorate some Down syndrome-associated phenotypes, chromosome 21 does not encode any components of the canonical SHH pathway. Here, we screened 163 chromosome 21 cDNAs in a series of SHH-responsive cell lines to identify chromosome 21 genes that modulate SHH signaling. We confirmed overexpression of trisomic candidate genes using RNA-seq in Ts65Dn and TcMAC21 cerebellum. Our study indicates that some chromosome 21 genes, including *DYRK1A*, upregulate SHH signaling while others, such as *HMGN1* and *MIS18A*, inhibit SHH signaling. Overexpression of genes involved in chromatin structure and mitosis, but not genes previously implicated in ciliogenesis, regulate the SHH pathway. Our data suggest that cerebellar hypoplasia and other phenotypes related to aberrant SHH signaling arise from the net effect of trisomy for multiple chromosome 21 genes rather than the overexpression of a single trisomic gene. Identifying which chromosome 21 genes modulate SHH signaling may suggest new therapeutic avenues for ameliorating Down syndrome phenotypes.

**One Sentence Summary:** Multiple chromosome 21 genes modulate Sonic hedgehog signaling, which is dysregulated in Down syndrome.

## INTRODUCTION

Down syndrome is a genetically complex condition with trisomy for >200 protein-coding genes contributing to an increased risk of more than 30 phenotypes (*1–3*). Although most Down syndrome-associated phenotypes remain unexplained at the molecular level, dysregulation of the Sonic hedgehog (SHH) signaling pathway may contribute to multiple Down syndrome phenotypes, including cerebellar hypoplasia, hippocampal learning and memory deficits, congenital heart defects, and more (*4*). Targeting the pleiotropic effects of aberrant SHH signaling is an attractive therapeutic strategy because a single treatment could theoretically rescue multiple phenotypes in individuals with Down syndrome. Given that there are currently no FDA-approved drugs to treat intellectual disability in Down syndrome, understanding the molecular mechanisms of SHH dysregulation during neurodevelopment is an aim with direct clinical relevance.

Cerebellar hypoplasia is one of a handful of phenotypes that occur in every individual with trisomy 21 and was the first Down syndrome-associated phenotype linked to abnormal SHH signaling (*5*). As measured by MRI, adults with Down syndrome have a disproportionally small cerebellum, even when adjusted for total brain volume (*6*). Ts65Dn mice, the most widely studied mouse model of Down syndrome, show cerebellar hypoplasia and a reduced density of cerebellar granule cell neurons; this reduced granule cell neuron density correctly predicted a similar deficit in people with Down syndrome (*7*).

We traced this reduction in granule cell neuron density to a defect in proliferation of granule cell precursors in early postnatal development (*5*). During normal development of the cerebellum, SHH acts as a mitogen for granule cell precursors (*8, 9*). Trisomic granule cell precursors isolated from postnatal day 6 (P6) Ts65Dn pups proliferate less in response to SHH than euploid cells, resulting in a small and hypocellular cerebellum (*5*). Neural crest cells isolated from the first pharyngeal arch of Ts65Dn embryos exhibit a parallel deficit in response to SHH, which acts as a mitogen for cells that will contribute to the mid and lower face (*10*). These experiments suggest that trisomic cells possess an intrinsic deficit in response to SHH signaling that is relevant to both cerebellar hypoplasia and craniofacial dysmorphology. SHH signaling is also required for normal development of the heart and enteric nervous system, supporting the hypothesis that abnormal signaling underlies increased risk of congenital heart defects and Hirschsprung disease in individuals with Down syndrome (*4*).

Consistent with this hypothesis, a single treatment with the SHH pathway agonist SAG on the day of birth rescues adult cerebellar morphology, performance on the Morris water maze, and some aspects of hippocampal long-term potentiation in Ts65Dn mice (*11*). Similarly, overexpression of a SHH transgene in the forebrain improves learning and memory in Ts65Dn mice, whereas overexpression of this SHH transgene in cerebellar Purkinje cells increases cerebellar volume but does not improve performance on learning and memory tasks (*12*). Another recent study showed that increasing the concentration of SAG normalizes expression of *OLIG2*, a chromosome 21 gene critical for oligodendrocyte development, in “brain-like” neural progenitors derived from human trisomy 21 iPSCs (*13*). Although these findings suggest that human trisomy 21 cells and mouse models of Down syndrome share a common defect in SHH signaling, treating humans with SAG or other SHH agonists may have unintended consequences on development. SHH signaling is a central developmental pathway involved in diverse processes from axis formation in early embryos to maintenance of stem cell niches in adults (*14*). Activating mutations in the SHH pathway are associated with medulloblastoma and basal cell carcinoma (*15–18*). SAG treatment also causes dose-dependent changes in cranial shape and size, which indicates that stimulation of SHH may have unwanted effects on skeletal development (*19*).

Given these limitations, targeting the trisomic genes responsible for abnormal SHH signaling may represent a better therapeutic strategy than activating the SHH pathway directly. Chromosome 21 does not encode known components of the SHH signaling pathway, and previous attempts to identify chromosome 21 genes involved in SHH signaling have focused on a small subset of candidate genes. The DYRK1A protein kinase has been identified as a modulator of SHH signaling, but returning *Dyrk1a* to disomy was not sufficient to rescue cerebellar volume in Ts65Dn mice (*20–22*). Overexpression of pericentrin (*PCNT*) was reported to disrupt ciliogenesis, which is required for canonical SHH signaling, but the mouse ortholog of *PCNT* is not trisomic in many of the mouse models with cerebellar hypoplasia, indicating that other trisomic genes are sufficient to cause this phenotype (*23*). Triplication of *APP* has also been proposed to inhibit SHH signaling by upregulating *PTCH1* (*24*).

In contrast to these candidate-based approaches, we sought to identify the chromosome 21 genes underlying disruption of SHH signaling using first principles and synthesis of available datasets. We propose that 1) Causal genes should be trisomic in mouse models with cerebellar hypoplasia; 2) Variation in causal genes may be linked to SHH phenotypes outside of the context of Down syndrome; 3) In the absence of genetic interactions, causal genes should inhibit SHH signaling when overexpressed; and 4) Causal genes should be expressed in the relevant cell types and misexpressed in trisomic cells. Here, we integrate data about cerebellar phenotypes collected in mouse models of Down syndrome, Mendelian disorders, a series of in vitro cDNA screens, and RNA-seq to show that the overexpression of multiple chromosome 21 genes modulates SHH signaling. Our findings prioritize four chromosome 21 genes (*B3GALT5*, *ETS2*, *HMGN1*, and *MIS18A*) that are trisomic in Ts65Dn mice, expressed in granule cell precursors, and inhibit proliferation when overexpressed in primary granule cell precursors.

## RESULTS

### Comparison of cerebellar phenotypes in Down syndrome mouse models

If a single trisomic gene is sufficient to cause a specific phenotype, individuals with trisomy for that gene will display the phenotype. In humans, this principle has been used to attempt to identify regions associated with intellectual disability, congenital heart anomalies, and other – mostly incompletely penetrant – aspects of the syndrome in rare individuals with partial trisomy 21 (*25, 26*). However, regional brain volume measurements are not available for human subjects with partial trisomy. We instead compared previously reported cerebellar volume or midline cross-sectional area measurements among mouse models at dosage imbalance for different subsets of chromosome 21 genes or their mouse orthologs (Fig. 1A **and table S1**). Cerebellar volumes ranged from 78% of euploid in Ts1Cje mice to 116% in 152F7 mice.

**Fig. 1.**
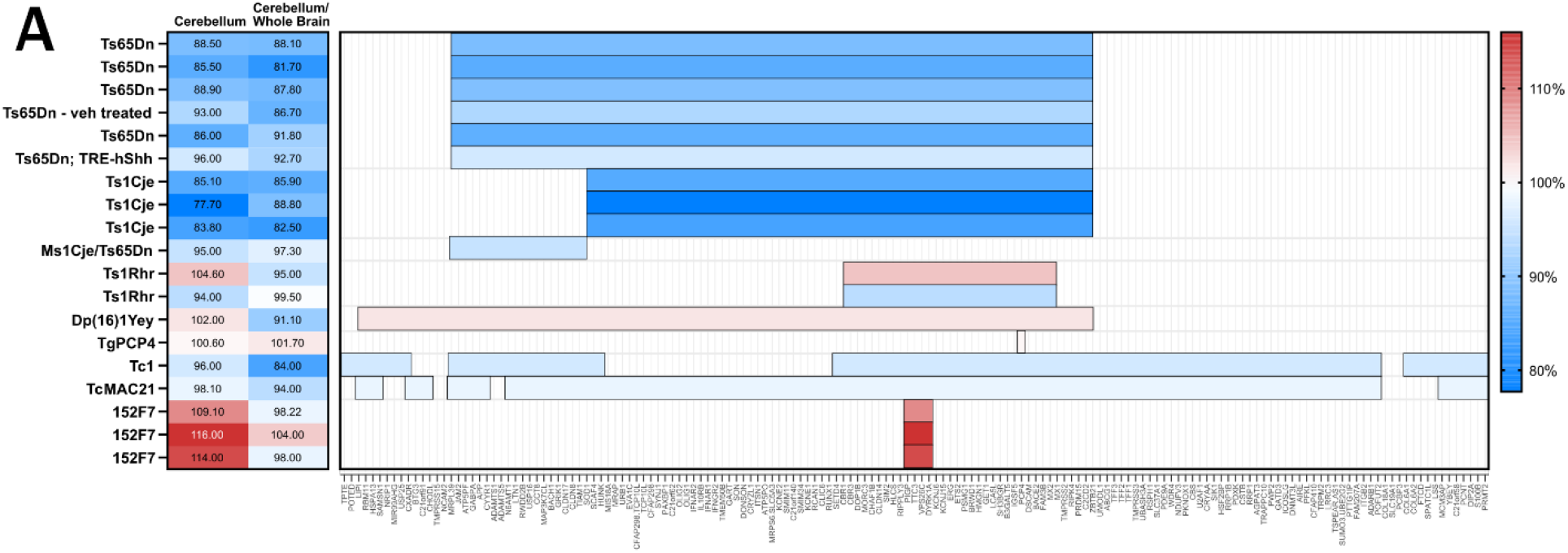
Comparison of cerebellar phenotypes in Down syndrome mouse models. **(A)** Previously published cerebellar volume/cross-sectional area and cerebellar volume/cross-sectional area normalized to whole brain are reported as a percent of euploid. Horizontal bars represent the chromosome 21 or mouse orthologous regions that are trisomic in each model. Color reflects the extent of cerebellar hypoplasia, where blue is most affected and red is least affected. Several additional studies quantifying cerebellar hypoplasia are generally consistent with these results but do not report cerebellar volume/cross-sectional area and cerebellar volume/cross-sectional area normalized to whole brain as a percent of euploid (*21, 30, 31*).

### Manual annotation of chromosome 21 genes related to SHH and ciliopathies

Disruption of the SHH pathway causes a range of well-characterized phenotypes, including holoprosencephaly, cerebellar hypoplasia, heart defects, skeletal abnormalities, and cancers such as medulloblastoma and basal cell carcinoma. To further understand how overexpression of chromosome 21 genes could affect SHH signaling, we manually annotated chromosome 21 genes associated with hedgehog-related phenotypes through a literature search, the Online Mendelian Inheritance in Man (OMIM) and Mouse Genome Informatics (MGI) databases, and the ciliary/centrosome database Cildb v3.0 (**table S2**). Of the 44 chromosome 21 genes with associated relevant phenotypes in OMIM, four genes (*CFAP298*, *CFAP410*, *PCNT*, and *RSPH1*) encode proteins involved in ciliogenesis. Mutations in an additional 12 genes (*CSTB*, *DSCAM*, *JAM2*, *KCNJ6*, *OLIG1*, *OLIG2*, *PRDM15*, *PSMG1*, *SOD1, SON*, *TRAPPC10*, and *WDR4*) are associated with cerebellar phenotypes or holoprosencephaly in humans or in mouse models. Unsurprisingly, many chromosome 21 genes have been reported to act upstream or downstream of SHH signaling in various cell types, including *ABCG1*, *ADARB1*, *APP*, *DYRK1A*, *GABPA*, *OLIG1*, *OLIG2*, *RUNX1*, *SIM2*, *SOD1*, *TIAM1*, and *USP25*. For several genes, multiple lines of evidence link their encoded proteins with SHH signaling or ciliogenesis. For example, mutant *Trappc10* mice possess septal defects, holoprosencephaly, anophthalmia, thymus hypoplasia, and cleft palate. The human *TRAPPC10* gene is located in a previously identified “holoprosencephaly critical region,” and knockdown of *TRAPPC10* impairs cilia formation in RPE cells (*27, 28*). Together, these annotations suggest that proteins encoded by chromosome 21 genes are components of both motile and primary cilia and may modulate SHH signaling through non-canonical pathways in diverse tissue types.

### Primary screen for chromosome 21 cDNAs that affect SHH signaling

Although several chromosome 21 genes have previously been associated with SHH signaling, most annotations derive from loss-of-function mutations rather than overexpression. To identify genes whose overexpression is sufficient to modulate SHH signaling, we designed a multilevel screen in zebrafish (*29*) and in four SHH-responsive cell types (Fig. 2A). We first screened a library of 163 human chromosome 21 cDNAs selected for high homology to mouse genes (**table S3**) in two well-established SHH-responsive cell lines: Shh-LIGHT2 cells, which express firefly luciferase from the SHH-responsive promoter of Gli1 (Fluc; 8xGliBS-FL) and renilla luciferase from a constitutive promoter (Rluc; pRL-TK, Promega), and SmoA1-LIGHT cells, which are based on Shh-LIGHT2 cells but also possess an oncogenic mutation in Smo (W539L) that activates SHH signaling in the absence of pharmacological stimulation (*15*).

**Fig. 2.**
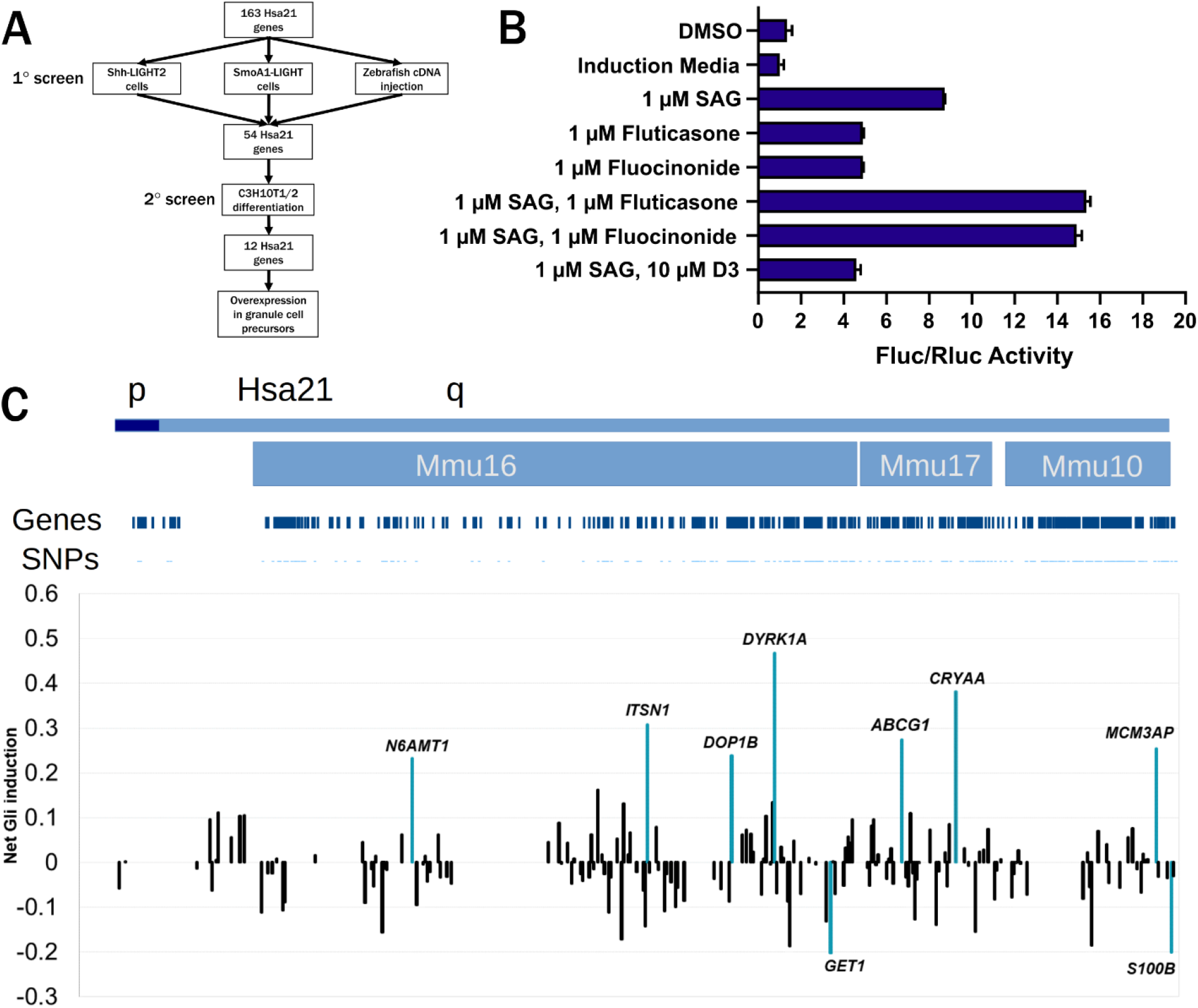
Overexpression of chromosome 21 cDNAs in Shh-LIGHT2 cells. **(A)** Screening strategy for chromosome 21 cDNAs in Shh-LIGHT2 and SmoA1-LIGHT cell lines, zebrafish embryos, C3H10T1/2 mesenchymal stem cell line, and primary granule cell precursors. **(B)** Fluc/Rluc activity in Shh-LIGHT2 cells exposed to SAG, the glucocorticoids fluocinonide and fluticasone, and vitamin D3 normalized to induction media control (n=2 independent experiments with 12 technical replicates per treatment). All graphs show mean ± SD unless otherwise noted. **(C)** Shh-LIGHT2 cells transfected with expression constructs for 163 chromosome 21 cDNAs and treated with SAG to induce SHH signaling (≥ 8 technical replicates per cDNA; see table S4 for wells per cDNA). Averaged Fluc/Rluc activity for each gene across the Shh-LIGHT2 screen was scaled to 0 to show signal deflections from baseline. Values less than zero represent loci that decrease SAG-induced activation of the SHH signaling pathway. The net activity of the 8xGliBS reporter for each cDNA is plotted in chromosomal order according to the sequence along the proximal-distal length of chromosome 21. Orthologous regions on mouse chromosomes 16, 17, and 10 are provided for additional context. Labeled cDNAs increased or decreased Fluc/Rluc activity by more than two SD.

Shh-LIGHT2 cells responded robustly to the hedgehog agonists fluocinonide, fluticasone, and SAG, whereas vitamin D3 inhibited SAG-induced reporter activity (Fig. 2B). Transient overexpression of nine genes increased or decreased the ratio of Fluc to Rluc activity by more than two standard deviations (z ≤ −2 or z ≥ 2) in Shh-LIGHT2 cells treated with SAG (**table S4**). Overexpression of *ABCG1*, *CRYAA*, *DOP1B*, *DYRK1A*, *ITSN1*, *MCM3AP*, and *N6AMT1* activated SHH signaling, whereas overexpression of *GET1* and *S100B* inhibited SHH (Fig. 2C).

In SmoA1-LIGHT cells, overexpression of *DYRK1A*, *IFNAR2*, and *MRPL39* increased SHH signaling by more than two standard deviations, and overexpression of *ABCG1*, *KCNE1*, *NDUFV3*, and *PRMT2* inhibited SHH signaling (Fig. 3A and **table S5**). We also identified an additional six genes that modulated SHH signaling by more than one standard deviation in both screens: *CHODL*, *HMGN1*, *KCNJ15*, *TTC3*, *UBASH3A*, and *VPS26C*. Of the twenty total genes identified in Shh-LIGHT2 or SmoA1-LIGHT screens, sixteen affected SHH signaling in the same direction in both cell lines: overexpression of *GET1*, *HMGN1*, *KCNE1*, *KCNJ15*, *NDUFV3*, *PRMT2*, and *UBASH3A* inhibited SHH signaling, overexpression of *CRYAA*, *DYRK1A*, *IFNAR2*, *ITSN1*, *MCM3AP*, *MRPL39*, *N6AMT1*, *TTC3*, and *VSP26C* upregulated SHH signaling, and *ABCG1*, *CHODL*, *DOP1B*, and *S100B* showed discordant directions of effect in the two cell lines (Fig. 3B). We previously screened the chromosome 21 cDNA library in developing zebrafish and identified eleven genes that caused gross morphological defects or lethality when overexpressed; seven of these genes affected development of structures that are substantially influenced by or dependent on SHH signaling (*29*). However, there was no overlap between any of these eleven genes and the twenty genes prioritized by the luciferase assays (Fig. 3C).

**Fig. 3.**
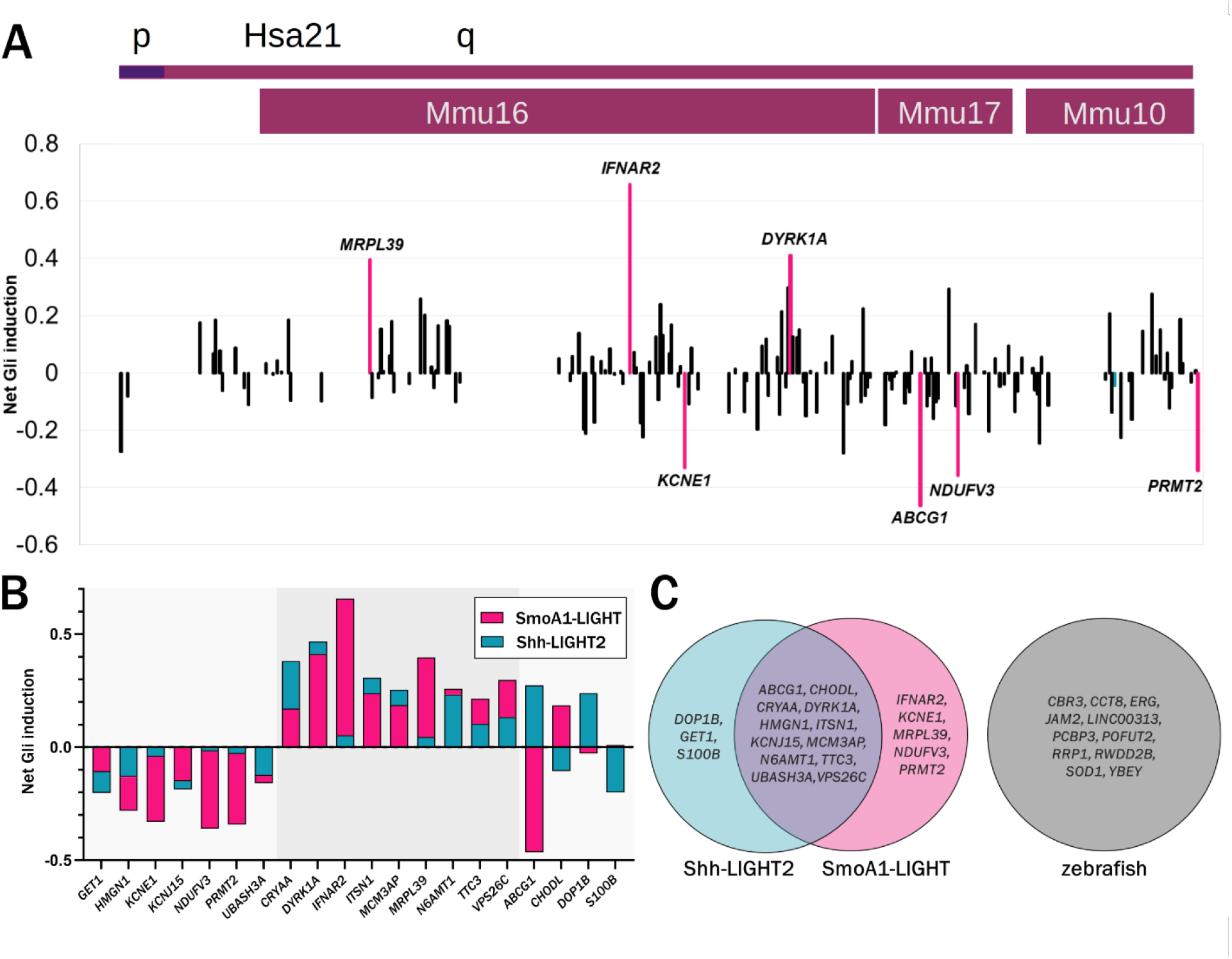
Overexpression of chromosome 21 cDNAs in SmoA1-LIGHT cells. **(A)** SmoA1-LIGHT cells transfected with expression constructs for 163 chromosome 21 cDNAs (≥ 8 technical replicates per cDNA; see table S5 for wells per cDNA). Averaged Fluc/Rluc activity for each gene across the SmoA1-LIGHT screen was scaled to 0 to show signal deflections from baseline. Labeled cDNAs increased or decreased Fluc/Rluc activity by more than two SD. **(B)** Comparison of net reporter induction after overexpression of twenty cDNAs identified in SmoA-LIGHT and Shh-LIGHT2 screens. Sixteen cDNAs have the same direction of effect in both screens, whereas four cDNAs have opposite directions of effect. **(C)** Comparison of cDNAs identified in two luciferase assays and a previous screen in developing zebrafish embryos (*29*).

We compared the results of our cDNA overexpression screens to four previously reported genome-wide siRNA knockdown and CRISPR knockout screens in 3T3-derived cell lines containing the 8xGliBS reporter (fig. S1 **and table S6**). Neither Shh-LIGHT2 nor SmoA1-LIGHT screens showed a significant correlation with two siRNA screens in NIH-3T3-ShhFL cells, which produce SHH endogenously (*32*). However, our Shh-LIGHT2 screen showed a weak negative correlation with two CRISPR knockout screens in 3T3 cells treated with ShhN, suggesting that knockout and overexpression of some chromosome 21 genes may have opposing effects on SHH signaling (*33, 34*). Of the 31 candidate genes identified by our cDNA and zebrafish screens, *DYRK1A, GET1, MCM3AP, PCBP3,* and *POFUT2* were identified in one or more of the four knockdown/knockout screens.

### Secondary screen using a functional cell-based assay of osteoblast differentiation

Based on our primary screen, we selected 54 chromosome 21 genes to further characterize in a functional cell-based assay. The C3H10T1/2 mesenchymal stem cell line undergoes SHH-dependent differentiation into osteoblasts and has been used to identify agonists and antagonists of the SHH signaling pathway (*35–37*). We transfected C3H10T1/2 cells with candidate cDNAs and quantified alkaline phosphatase activity, an early marker of osteoblast differentiation. In the absence of SAG treatment, overexpression of *GLI1* was sufficient to induce osteoblast differentiation (Fig. 4A). Stimulation of osteoblast differentiation by 200 nM SAG was inhibited by co-treatment with 2 uM cyclopamine and by overexpression of the heterotrimeric G-protein subunit GαS (*GNAS*), which inhibits SHH signaling via protein kinase A (PKA) (*33, 38*). Overexpression of the previously identified regulator of SHH signaling *MOSMO* had no effect on alkaline phosphatase activity, whereas overexpression of *GLI1* further induced osteoblast differentiation even in the presence of SAG (*33*).

**Fig. 4.**
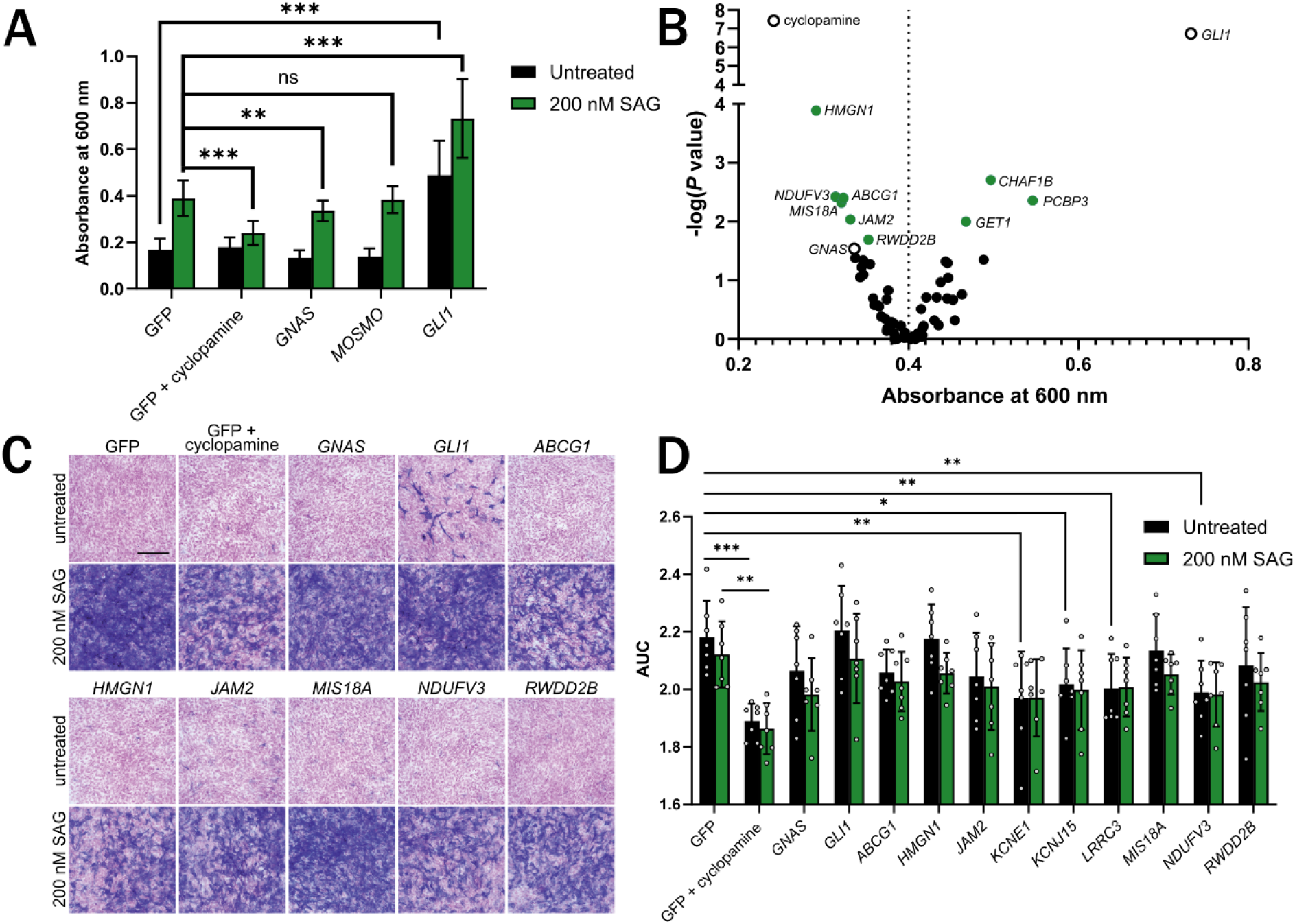
Overexpression of chromosome 21 genes affects osteoblast differentiation of C3H10T1/2 cells. **(A)** Overexpression of *GLI1* promotes osteoblast differentiation in the presence or absence of SAG, whereas treatment with cyclopamine or overexpression of *GNAS* inhibits SAG-induced osteoblast differentiation (n=20). ***P<0.001, **P<0.01, *P<0.05 (two-tailed unpaired Student’s t-test). **(B)** Quantification of alkaline phosphatase activity in C3H10T1/2 cells transfected with chromosome 21 cDNAs and treated with SAG (n=20). Multiple comparisons were corrected for by controlling the false discovery rate; green circles denote cDNAs with q<0.1. Open circles denote controls (Kruskal-Wallis test followed by Dunn’s post-hoc test). **(C)** Representative alkaline phosphatase staining in C3H10T1/2 cells transfected with chromosome 21 cDNAs and counterstained with nuclear fast red (n=3). Scale bar: 100 μm. **(D)** MTT viability assay in C3H10T1/2 cells transfected with chromosome 21 cDNAs (n=7). Y-axis represents area under the curve (AUC) of cell viability 48, 72, and 96 hours after transfection (two-way ANOVA followed by Fisher’s LSD test). Differences reported as statistically significant have q<0.05.

In C3H10T1/2 cells treated with SAG, overexpression of six chromosome 21 cDNAs (*ABCG1*, *HMGN1*, *JAM2*, *MIS18A*, *NDUFV3*, and *RWDD2B*) significantly reduced osteoblast differentiation compared to control, indicating that overexpression of these cDNAs attenuated SHH signaling (Fig. 4B and fig. S2). Overexpression of three chromosome 21 cDNAs (*CHAF1B*, *GET1*, and *PCBP3*) significantly increased osteoblast differentiation compared to control. Staining of cells for alkaline phosphatase activity in a subset of cDNAs confirmed inhibition of osteoblast differentiation and suggested a possible reduction in cell density following transfection of some cDNAs (Fig. 4C). Because reduced viability could affect osteoblast differentiation independently of SHH signaling, we assessed cell viability at three time points post-transfection using a MTT assay. Both cDNA (f(12, 156)=5.327, P<0.0001) and SAG treatment (f(1, 156) = 6.474, P=0.0119) had a significant effect on viability, but the interaction between these terms was not significant (Fig. 4D and fig. S2). In untreated cells, overexpression of *KCNE1*, *KCNJ15*, *LRRC3*, and *NDUFV3* and treatment with cyclopamine reduced cell viability compared to control. In cells treated with SAG, only cyclopamine treatment significantly affected viability.

### Expression of candidate genes in developing cerebellum

To determine whether candidate genes are expressed in a SHH-responsive tissue relevant to Down syndrome-associated cerebellar hypoplasia, we performed RNA-seq on P6 cerebellum collected from Ts65Dn (n=4 trisomic and 4 euploid littermates) and TcMAC21 (n=4 trisomic and 4 euploid littermates) pups. At this stage of development, the cerebellum is composed primarily of proliferating granule cell precursors and differentiating granule cells (*39, 40*). We previously found that granule cell precursors isolated from P6 Ts65Dn pups respond less to the mitogenic effects of SHH than euploid cells, and by P6, cerebellar cross-sectional area is significantly reduced in Ts65Dn (*5*). For TcMAC21 samples, length-normalized counts for human chromosome 21 transcripts were added to counts for corresponding mouse orthologs and compared to euploid counts. Trisomic genes were overexpressed by an average of 1.45±0.29 in Ts65Dn mice and 1.81±1.18 in TcMAC21 mice compared to euploid (Fig. 5A). The majority of trisomic genes with detectable expression in Ts65Dn mice had fold changes between 1.3 and 1.7, whereas TcMAC21 samples had a higher proportion of trisomic genes with fold changes above 1.7 (Fig. 5B). Arranged by chromosomal position, expression patterns were consistent with the previously reported breakpoint of the Ts65Dn 17^16^ chromosome and the four deletions reported in the TcMAC21 HSA21q-MAC hybrid chromosome (Fig. 5C) (*41, 42*). Expression of human chromosome 21 genes in TcMAC21 cerebellum was positively correlated with previously published P1 forebrain expression levels (r=0.39, P=2.3e-05) (Fig. 5D), and 31 human genes were not detected in TcMAC21 P1 forebrain or P6 cerebellum (Fig. 5E).

**Fig. 5.**
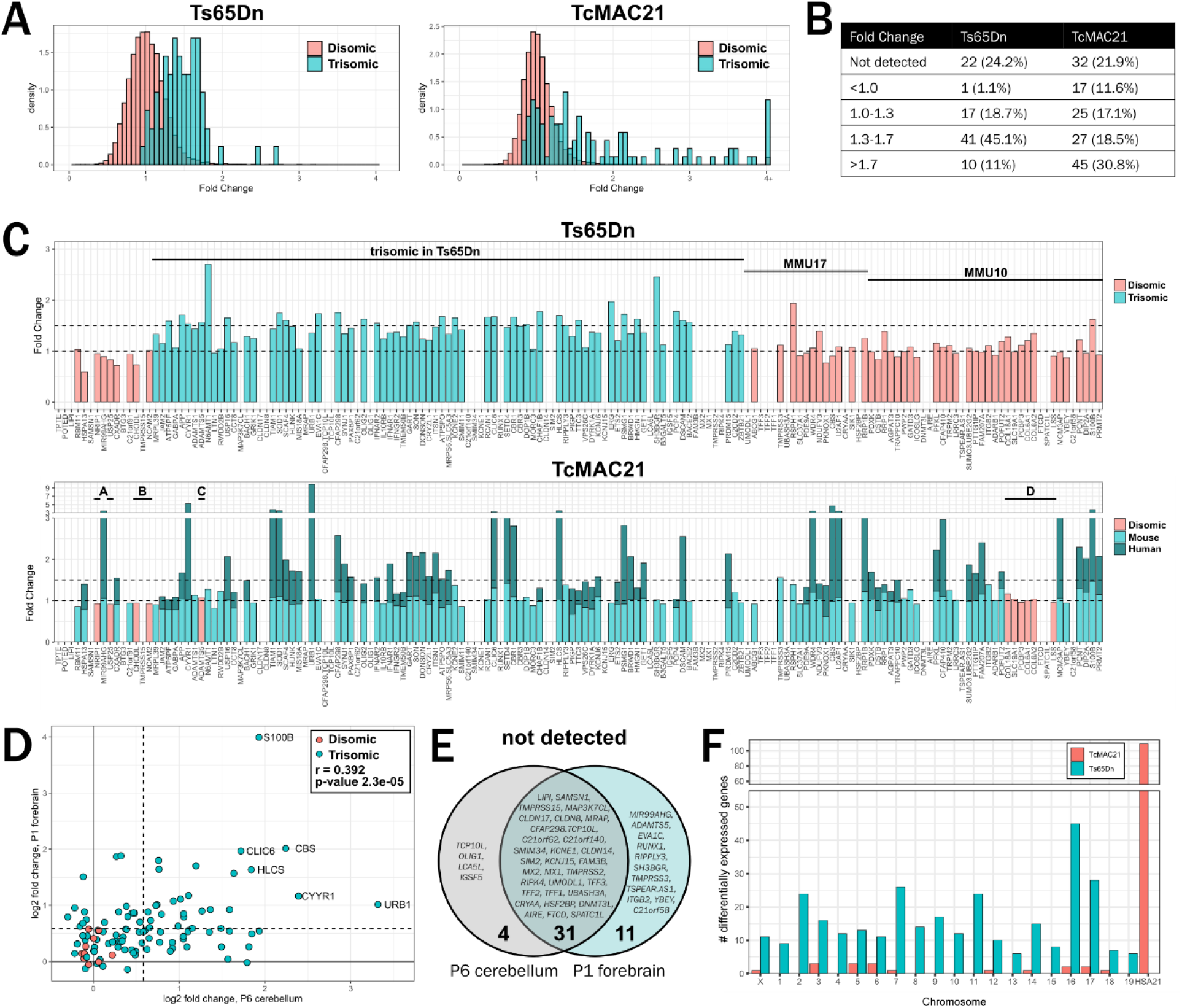
Expression pattern of chromosome 21 genes and their mouse orthologs in Ts65Dn and TcMAC21 cerebellum. **(A)** Density histograms of disomic (salmon) and trisomic (teal) fold changes in Ts65Dn and TcMAC21 cerebellum (n=4 Ts65Dn, 4 Ts65Dn euploid littermates, 4 TcMAC21, 4 TcMAC21 euploid littermates). Plot represents 13,807 detectable transcripts. **(B)** Trisomic gene fold changes binned by expression levels. **(C)** Fold changes of chromosome 21 genes and their mouse orthologs arranged in chromosomal order from proximal to distal. Chromosome 21 orthologs are located on mouse chromosome 16 (MMU16), MMU17, and MMU10. For TcMAC21, teal represents the proportion of length-normalized reads contributed by mouse copies and dark teal represents reads derived from the human chromosome. Four previously reported deletions are labeled A through D. Five human genes that were detected in TcMAC21 but have no expression of mouse orthologs for normalization (*POTED*, *BTG3*, *RUNX1*, *C21orf58*, and *TSPEAR.AS1*) are excluded. **(D)** Scatterplot of chromosome 21 log2 fold changes in TcMAC21 P6 cerebellum and P1 forebrain (*42*). Pearson correlation coefficient r=0.392 and P=2.3e-05. **(E)** Chromosome 21 transcripts not detected in P6 cerebellum, P1 forebrain, or both. **(F)** Chromosomal locations of differentially expressed genes in Ts65Dn and TcMAC21 cerebellum. Trisomic genes are located on MMU16 and MMU17 in Ts65Dn mice and HSA21 in TcMAC21 mice.

We also identified differential expression of disomic genes in both Ts65Dn and TcMAC21 models (Fig. 5F, fig. S3**, table S7, and table S8**). Although expression levels in Ts65Dn and TcMAC21 cerebella were positively correlated (r=0.529 and P=2.2e-16), only two disomic genes, *Lrch4* and *Snhg11*, were significantly differentially expressed in both models using a false discovery rate of 0.05 (fig. S3). Gene ontology and gene set enrichment analyses of differentially expressed genes in Ts65Dn samples suggested changes in gene expression related to nervous system development, higher mental function, and cholesterol biosynthesis (fig. S4 **and** S5 **and table S9**). Ts65Dn samples also showed reduced expression of mitotic and cell cycle pathways and increased expression of genes related to protein translation initiation and elongation (fig. S5). The trisomic chromatin modifiers and remodelers *Chaf1b*, *Hmgn1*, *Setd4*, and *Brwd1* were significantly upregulated, and non-trisomic epigenetic regulators *Rps6ka5*, *Rere*, *Brd4*, *Kdm7a*, and *Top2a* were dysregulated (fig. S6). Elements of the Polycomb repressive complex (*Mbd6*, *Pcgf2*, and *Auts2*) and SWI/SNF complex (*Arid1a*, *Arid1b*, and *Bicra*) were also upregulated.

### Synthesis of expression and SHH screen data to prioritize candidate genes

We next integrated expression data with our primary and secondary SHH screen data. Leading candidate genes should be expressed in developing cerebellum, trisomic in mouse models with cerebellar hypoplasia, and consistently inhibit SHH across in vitro screens. Eighteen genes were not detected in our RNA-seq data, had FPKM (fragments per kilobase of exon per million mapped) values <1 in 13 human cerebellar samples ranging from 12 post-conception weeks to 4 months (BrainSpan Atlas of the Developing Human Brain), and had TPM (transcripts per million) <1 in mouse P2 and P11 granule cell precursor and granule cell populations (Fig. 6A and **table S10**) (*39, 43*). Although these 18 genes may contribute to dysregulated SHH signaling in other tissues, such as the heart or craniofacial skeleton, they appear as unlikely candidates for cerebellar hypoplasia.

**Fig. 6.**
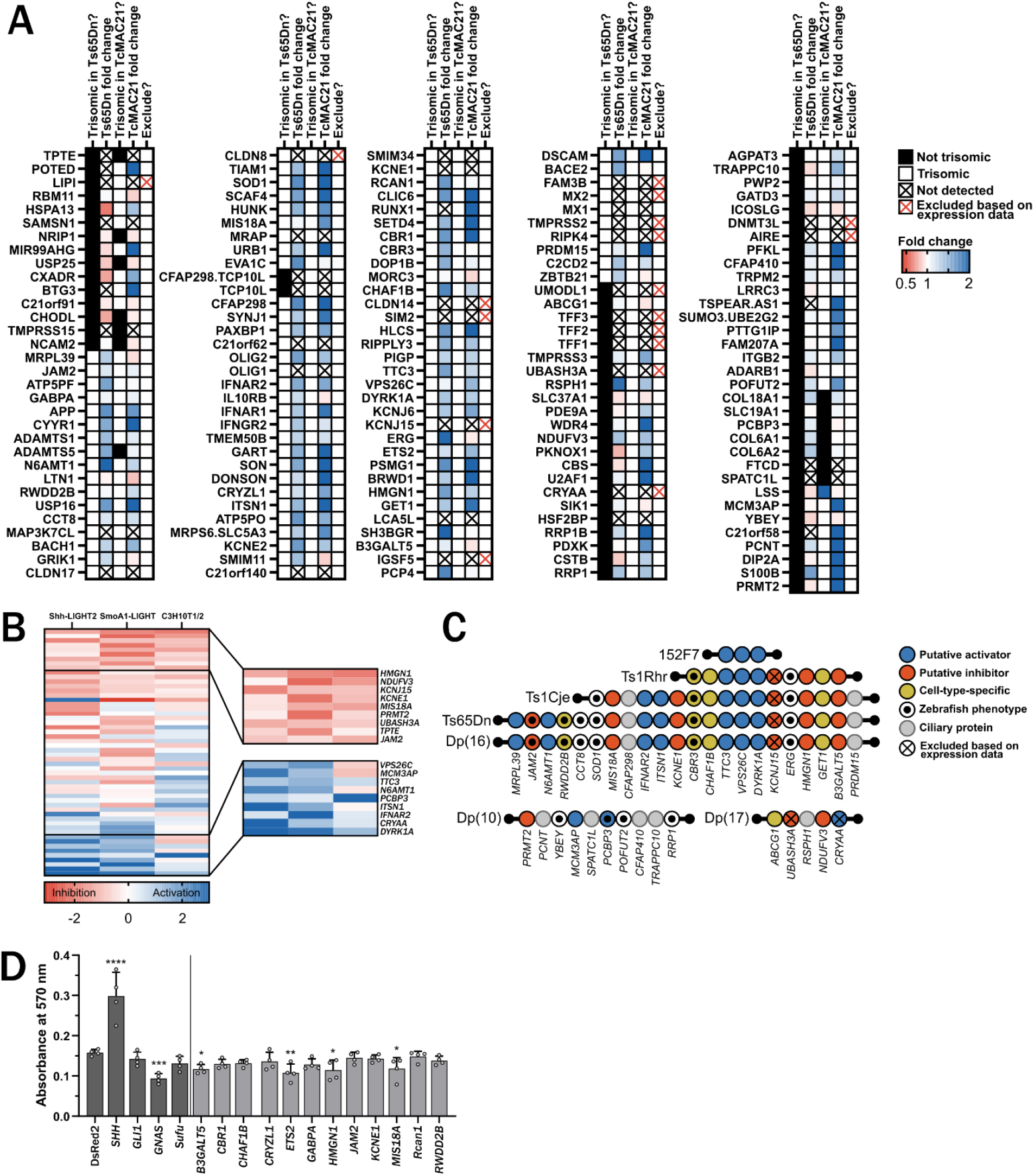
Prioritization of candidate cDNAs and overexpression in primary granule cell precursors. **(A)** Summary of expression data in developing cerebellum. Black boxes indicate genes that are not trisomic in Ts65Dn and TcMAC21 mouse models, and white boxes indicate genes that are trisomic in these models. Fold change is indicated by color with red signifying decreased expression and blue signifying increased expression. Transcripts with black crosses were not detected in our RNA-seq dataset, and transcripts with red crosses were excluded based on our expression data, expression in the BrainSpan Atlas of the Developing Human Brain, and single cell RNA-seq data from euploid mouse granule cell precursors and granule cells. **(B)** Comparison of the effects of 54 chromosome 21 cDNAs in Shh-LIGHT2, SmoA1-LIGHT, and C3H10T1/2 screens. cDNAs are sorted by average z-score with red signifying inhibition and blue signifying activation of the SHH pathway. Inset shows top and bottom ranked cDNAs. **(C)** Chromosomal locations of the mouse orthologs of candidate cDNAs in Down syndrome mouse models. *LINC00313* (*C21ORF84*) and *TPTE* are not shown. *LINC00313*, which was identified in the zebrafish screen, is a human-specific gene and not present in the listed mouse models. *TPTE* is located on the short arm of human chromosome 21 and has a putative homolog on mouse chromosome 8. **(D)** Lentiviral overexpression of candidate genes inhibits proliferation of granule cell precursors treated with 6 nM SAG and pulsed with EdU for 24 hours (n=4). ****P <0.0001, ***P<0.001, **P<0.01, *P<0.05 (one-way ANOVA followed by Fisher’s LSD test). Differences reported as statistically significant have q<0.05.

Synthesis of data from our primary luciferase screens and secondary C3H10T1/2 screen also revealed candidate genes with the most consistent effects across cell lines (Fig. 6B). For example, overexpression of *HMGN1* consistently inhibited SHH, whereas overexpression of *DYRK1A* consistently activated SHH. Most cDNAs showed relatively consistent effects across cell types, but some cDNAs, such as *ABCG1* and *GET1*, showed strong but discordant effects across screens. We mapped candidate cDNAs to mouse models and identified a subset of six genes that appear to inhibit SHH across screens and have mouse orthologs located on mouse chromosome 16 (Fig. 6C). Putative activators of SHH that are trisomic in Ts65Dn mice (*DYRK1A*, *IFNAR2*, *ITSN1*, *MRPL39*, *N6AMT1*, *TTC3*, and *VPS26C*) may provide compensatory effects, while putative inhibitors of SHH with mouse orthologs on chromosomes 10 and 17 (*NDUFV3*, *PRMT2*, and *UBASH3A*) may inhibit SHH via a mechanism independent of dysregulated SHH signaling in Ts65Dn cells.

### Overexpression of four candidate genes inhibits proliferation of primary granule cell precursors

To evaluate top candidate cDNAs in a context relevant to cerebellar hypoplasia, we cloned 12 chromosome 21 cDNAs into lentiviral vectors, overexpressed them in primary euploid granule cell precursors, and quantified proliferation via incorporation of EdU. As expected, overexpression of *SHH* itself significantly increased proliferation, and overexpression of *GNAS*, which has been identified as a tumor suppressor gene in the SHH subtype of medulloblastoma, significantly inhibited proliferation (Fig. 6D) (*44*). Of the 12 chromosome 21 cDNAs, overexpression of four (*B3GALT5*, *ETS2*, *HMGN1*, and *MIS18A*) significantly reduced proliferation compared to overexpression of DsRed2. These results suggest that at least some of the candidate cDNAs identified in the luciferase, zebrafish, and C3H10T1/2 screens also modulate SHH signaling in the developing cells of the cerebellum.

## DISCUSSION

Our data provide novel insights into the complex genetic architecture of aberrant SHH signaling in Down syndrome. We previously showed that a reduced mitogenic response to SHH underlies cerebellar hypoplasia in Ts65Dn mice but lacked a clear understanding of which trisomic genes contribute to this phenotype (*5*). In this study, we prioritized chromosome 21 genes that consistently modulate SHH signaling in a variety of cellular contexts and identified four genes (*B3GALT5*, *ETS2*, *HMGN1*, and *MIS18A*) that impair the proliferation of cerebellar granule cell precursors when individually overexpressed (fig. S7).

In contrast to previous hypothesis-driven approaches, our study provides quantitative data about the individual effects of nearly all chromosome 21 protein-coding genes conserved between human and mouse. While trisomy of any chromosome has the potential to impair proliferation via aneuploidy stress, our data show that overexpression of specific chromosome 21 genes inhibits the proliferation of granule cell precursors (*45*). In fact, overexpression of 127 of the 163 cDNAs had no effect in the luciferase, zebrafish, C3H10T1/2, or granule cell precursor assays, indicating that cDNA overexpression does not have a non-specific effect on SHH signaling and, barring genetic interactions, excludes these genes as candidates. Lack of effect in our SHH screens does not eliminate them as contributors to a general destabilization of the trisomic transcriptome, nor does it consider effects in the context of a transcriptome destabilized by trisomy for individually benign trisomic genes.

Our results provide evidence for why Down syndrome mouse models present with variable severities of cerebellar hypoplasia (Fig. 1). Ts65Dn mice and Ts1Cje mice possess a similar reduction of cerebellar volume normalized to total brain volume (*7, 11, 12, 46-49*). Ms1Cje/Ts65Dn mice do not show substantial cerebellar hypoplasia, although only three trisomic animals were analyzed (*46*). Ts1Rhr mice show more subtle cerebellar hypoplasia than Ts65Dn and Ts1Cje mice (*47, 50*). Comparing these four models suggests that at least one gene in the region that is trisomic in both Ts1Cje and Ts65Dn mice but is not trisomic in Ts1Rhr mice (*Sod1* to *Setd4* and *Ripk4* to *Zbtb21*) contributes to cerebellar hypoplasia. An additional gene or genes may contribute to the mild cerebellar hypoplasia observed in Ts1Rhr mice (*Cbr1* to *Fam243b*). 152F7 mice, which contain a YAC with *PIGP, TTC3, VPS26C,* and *DYRK1A*, show increased cerebellar volume relative to control, suggesting that overexpression of this region provides a compensatory effect (*51*). Although Tc1 mice also display cerebellar hypoplasia, interpreting the genetic contributions to this phenotype is challenging due to mosaicism and complex rearrangements in the Tc1 human chromosome (*52, 53*). Overexpression of *PCP4*, Purkinje cell protein 4, does not affect cerebellar volume (*54*).

The results from our SHH screen are consistent with a model in which *B3galt5*, *Ets2*, *Hmgn1*, and *Mis18a* contribute to the severe cerebellar hypoplasia observed in Ts65Dn and Ts1Cje mice, *Hmgn1* and *Ets2* contribute to the milder hypoplasia in Ts1Rhr mice, and *Ttc3*, *Vps26c*, and *Dyrk1a* provide a compensatory effect in Ts65Dn, Ts1Cje, Ts1Rhr, and 152F7 mice. Differences in gene content may also explain why relatively mild cerebellar hypoplasia was reported in Dp(16)1Yey and in TcMAC21 mice, despite these models containing more trisomic genes than either Ts65Dn or Ts1Cje (*42, 55*). For example, overexpression of *MCM3AP*, a putative activator of SHH signaling and MMU10 ortholog, could provide a compensatory effect in TcMAC21 cerebellum.

Interpreting the contributions of trisomic genes to cerebellar hypoplasia is further complicated by differences in expression of trisomic genes between models. TcMAC21 and Tc1 models rely on appropriate function of human DNA regulatory elements in mouse cells, and gene expression may differ between models with segmental duplications (e.g., Dp(16)1Yey and Ts1Rhr) versus freely segregating chromosomes (Ts65Dn) (*56*). For example, we found that *B3GALT5*, a putative inhibitor of SHH signaling, has a fold change of 0.92 in TcMAC21 despite this model having two copies of mouse *B3galt5* and one copy of human *B3GALT5.* Moreover, although our SHH screen and RNA-seq data support an oligogenic or polygenic explanation for cerebellar hypoplasia in mouse models, testing this hypothesis by returning candidate genes to disomy would be technically challenging due to difficult husbandry, relatively subtle phenotypes, and high interindividual variability (*57, 58*).

The molecular mechanisms by which chromosome 21 genes inhibit SHH signaling merit additional exploration. It is surprising that overexpression of several known ciliary genes (*CFAP298*, *CFAP410*, *RSPH1*, *SPATC1L*, and *TRAPPC10*) had no effect in our luciferase screens, consistent with a previous report that overexpression of *CFAP298*, *CFAP410*, and *TRAPPC10* did not alter cilia formation (*23*). Instead, we identified a number of regulators of mitosis and chromatin structure, including *CHAF1B*, *HMGN1*, *MCM3AP*, *MIS18A*, and *N6AMT1*, two involved in endocytosis, *ITSN1* and *VPS26C*, and a cholesterol transporter, *ABCG1*. These results suggest that rather than inhibiting the canonical SHH pathway directly, overexpression of some chromosome 21 genes may affect cell state, epigenetic regulation, and progression through the cell cycle. This hypothesis is supported by differential expression of chromatin regulators in Ts65Dn cerebellum and gene set enrichment analysis showing reduced expression of transcripts encoding mitotic proteins.

A promising candidate for disruption of normal chromatin structure is *HMGN1*, which was overexpressed in Ts65Dn cerebellum and inhibited SHH signaling when overexpressed in Shh-LIGHT2 and SmoA1-LIGHT cells, C3H10T1/2 cells, and primary granule cell precursors. In our previous zebrafish screen, only nine of 120 embryos survived injection of 50 pg *HMGN1* mRNA, and of the nine surviving embryos, four had missing melanocytes (*29*). However, this finding was not reproduced in a secondary screen. In *Xenopus laevis*, injection of HMGN1 protein causes body axis curvature, cyclopia, and microcephaly, which are all phenotypes associated with aberrant SHH signaling (*59*). *Hmgn1* is expressed in the pharyngeal arches of *Xenopus* embryos, and knockdown of *Hmgn1* disrupts cranial neural crest streams, resulting in hypoplastic craniofacial cartilage (*60*).

*HMGN1* encodes a non-histone chromosomal protein that competes for binding with histone H1 (*61*). Binding of HMGN1 reduces chromatin compaction and is associated with lineage-specific regulatory elements (*62*). HMGN1 expression levels are correlated with the transition from proliferation to differentiation in stem cells, and in primary rat osteoblasts, *Hmgn1* is preferentially expressed in proliferating cells with a decline in expression at the onset of mineralization (*63*). Loss of *Hmgn1* and *Hmgn2* in mouse embryonic fibroblasts increases the efficiency of reprogramming into iPSCs, suggesting that HMGN proteins help to stabilize cell identity (*62*). In B cells, *HMGN1* overexpression results in a loss of H3K27me3, a gain of H3K27ac, and a global increase in transcription (*64, 65*). HMGNs act upstream of *Olig1* and *Olig2* during oligodendrocyte differentiation, indicating a possible interplay between SHH signaling, *HMGN1*, and *OLIG1/2* during neurodevelopment (*13, 66*). Together, the known roles of HMGN1 suggest that HMGN1 overexpression could disrupt the proliferation of granule cell precursors by altering epigenetic marks, disrupting the balance of proliferation and differentiation (e.g., precocious differentiation), or promoting differentiation along an alternative cell state trajectory, such as differentiation into astrocytes (*67*).

Our screen focused on inhibitors of SHH activity, but several chromosome 21 cDNAs consistently activated SHH signaling across cell types. Therapeutic interventions targeting trisomic activators of SHH could worsen SHH-associated phenotypes in people with Down syndrome. In particular, DYRK1A stimulated SHH signaling in our luciferase screens and was previously reported to activate SHH by phosphorylating GLI1 and promoting its retention in the nucleus (*20, 68*). Overexpression of *DYRK1A* was previously reported to induce osteoblast differentiation of C3H10T1/2 cells (*68*), though this activation did not reach statistical significance in our screen. DYRK1A is commonly proposed as a target for treating Down syndrome-associated intellectual disability (*69, 70*), but we recommend monitoring potential worsening of phenotypes in SHH-responsive tissues, such as the cerebellum, heart, and bone, in preclinical studies of DYRK1A inhibitors (*71–75*).

Our screening paradigm made significant progress towards understanding how the overexpression of chromosome 21 genes influences SHH signaling. However, no cell culture methods can fully represent the complex effects of trisomy 21 on human development. Individual overexpression of cDNAs cannot reproduce the effects of simultaneous overexpression of chromosome 21 genes in the context of trisomy or detect genetic interactions between sets of trisomic genes. Our library contains most conserved chromosome 21 protein-coding genes but does not include several genes that may influence neurodevelopment, including *PCNT* and *SON* (*23, 76–78*). Transient transfection likely causes supraphysiological overexpression of cDNAs, and we did not comprehensively confirm expression of each cDNA, leading to possible false negatives. Lastly, overexpression of some cDNAs may cause lethality rather than inhibiting SHH directly. Overexpression of potassium channel subunits *KCNE1* and *KCNJ15* and the mitochondrial subunit *NDUFV3* inhibited SHH but also affected viability in C3H10T1/2 cells. Future work must confirm that candidate genes modulate SHH in vivo and at expression levels mirroring the expected ∼1.5-fold increase observed in trisomy.

Our study established *B3GALT5*, *ETS2*, *HMGN1*, and *MIS18A* as likely regulators of proliferation in the developing cerebellum. However, despite completing three parallel screens and a secondary screen, we lack a simplistic answer as to how trisomy 21 causes cerebellar hypoplasia in people with Down syndrome. Much research has devoted itself to identifying “the” chromosome 21 gene responsible for each Down syndrome-associated phenotype. Current methods, such as mapping panels and returning trisomic genes to disomy in the context of trisomy, are ill equipped to deal with complex genetic interactions and compensatory effects. Our findings suggest that for developmental phenotypes like intellectual disability, determining the individual effects of trisomic genes across the lifetime may require the development and application of new techniques and the reframing of Down syndrome as a complex genetic disorder.

## MATERIALS AND METHODS

### Animals

All procedures met the requirements of the National Institutes of Health Guide for the Care and Use of Laboratory Animals and were approved by and carried out in compliance with the Johns Hopkins University Animal Care and Use Committee. Founder B6EiC3H-a/A-Ts65Dn Stock No. 001924 (Ts65Dn) mice were obtained from The Jackson Laboratory and maintained as an advanced intercross on a C57BL/6J × C3H/HeJ genetic background. These mice represent the original Davisson strain (*79–81*). TcMAC21 mice were generated as previously described and maintained on a C57BL/6J (B6) × DBA/2J (D2) background (*42*). Ts65Dn mice were genotyped by PCR and TcMAC21 mice were genotyped by GFP fluorescence. For RNA-seq, cerebella from pairs of trisomic pups and euploid littermates were isolated from two (Ts65Dn) or three (TcMAC21) litters. Euploid pups for granule cell precursors were C57BL/6J × C3H/HeJ, and cerebella of both sexes were pooled within litters. All experiments were performed on postnatal day 6 (P6).

### Plasmids

Luciferase assays were carried out using the Hsa21 Gene Expression Set in the pCSDest2 vector (https://www.addgene.org/kits/reeves-hsa21-set/). cDNAs for the C3H10T1/2 differentiation assay were subcloned into the pcDNA™6.2/EmGFP-Bsd/V5-DEST vector (Invitrogen, V36620) and included full length cDNAs for KCNE1, DOP1B, and Rcan1, which were truncated in our original pCSDest2 cDNA library. cDNAs for lentiviral transduction were subcloned into the plenti-CAG-gate-FLAG-IRES-GFP vector. The plenti-CAG-gate-FLAG-IRES-GFP vector was a gift from William Kaelin (Addgene plasmid #107398; http://n2t.net/addgene:107398; RRID:Addgene_107398) (*82*). To facilitate efficient subcloning, the vector’s kanamycin resistance gene was replaced with the ampicillin resistance gene from the pcDNA™6.2/EmGFP-Bsd/V5-DEST vector by digesting both vectors with BspHI and ligating with T4 DNA ligase.

Unless otherwise noted, Hsa21 cDNAs were acquired from the Hsa21 Gene Expression Set as previously described and subcloned using Gateway cloning (*29*). JAM2 was obtained from the Hsa21 Gene Expression Set and subcloned using TOPO cloning. KCNE1 (ThermoFisher, Ultimate ORF Clone IOH54610) and TRAPPC10 (ThermoFisher, Ultimate ORF Clone IOH53207) in pENTR221 were obtained from Johns Hopkins University Hit Genomics Services. mRcan1 (Dharmacon, Mammalian Gene Collection 4236038) in pCMV-SPORT6 was subcloned using TOPO cloning. DOP1B (HsCD00431873) in pENTR223.1 was obtained from The ORFeome Collaboration and subcloned using TOPO cloning.

pENTR-DsRed2 N1 (CMB1) was a gift from Eric Campeau (Addgene plasmid #22523; http://n2t.net/addgene:22523; RRID:Addgene_22523). pDONR223_GLI1_WT was a gift from Jesse Boehm & William Hahn & David Root (Addgene plasmid #82123; http://n2t.net/addgene:82123; RRID:Addgene_82123) (*83*). pEGFPC3-mSufu was a gift from Aimin Liu (Addgene plasmid #65431; http://n2t.net/addgene:65431; RRID:Addgene_65431) and was subcloned using TOPO cloning (*84*). GNAS (HsCD00288799) in pENTR223 was obtained from The ORFeome Collaboration and subcloned using TOPO cloning. SHH (HsCD00082632) in pENTR223.1 was obtained from The ORFeome Collaboration. MOSMO (EX-H4481-M02) in pReceiver-M02 was obtained from GeneCopoeia. pMD2.G was a gift from Didier Trono (Addgene plasmid #12259; http://n2t.net/addgene:12259; RRID:Addgene_12259). psPAX2 was a gift from Didier Trono (Addgene plasmid #12260; http://n2t.net/addgene:12260; RRID:Addgene_12260). Plasmids for transfection were purified using endotoxin-free midiprep kits.

### Cell culture

Shh-LIGHT2 and SmoA1-LIGHT were gifts from Philip Beachy and colleagues and were derived from the original stocks created by this group at Johns Hopkins University (*15*). Shh-LIGHT2 and SmoA1-LIGHT cells were grown in Dulbecco’s Modified Eagle’s Medium (DMEM; Gibco, 11965092) supplemented with 10% calf serum (Sigma, C8056 or N4637) and 1% penicillin-streptomycin (Quality Biological, 50-751-7267). Shh-LIGHT2 cultures were kept under antibiotic selection with 400 μg/mL Geneticin (Gibco, 10131035) and 150 μg/mL Zeocin (Invitrogen, R25001) and SmoA1-LIGHT cells were cultured with 400 μg/mL Geneticin and 100 μg/ml Hygromycin B (Corning, 30-240-CR). C3H10T1/2 cells (ATCC, CCL-226) were maintained in DMEM supplemented with 10% fetal bovine serum (HyClone, SH30071.03), 2 mM L-Glutamine (Quality Biological, 118-084-721), and 1% penicillin-streptomycin. 293FT cells (Invitrogen, R70007) were maintained in DMEM with 10% FBS, 0.1 mM MEM Non-Essential Amino Acids (Gibco, 1114050), 6 mM L-glutamine, 1 mM MEM sodium pyruvate (Sigma, S8636), and 1% penicillin-streptomycin with 500 ug/mL Geneticin. Primary granule cell precursors were maintained in neurobasal medium (Gibco, 21103049) with 2 mM Glutamax (Gibco, 35050061), 1% penicillin-streptomycin, 1 mM sodium pyruvate (Sigma, S8636), and 2% B27 (Gibco, 12587010) with 6 nM SAG (Calbiochem, 566661).

### Luciferase reporter assays

To quantify hedgehog pathway activity in Shh-LIGHT2, cells were removed from antibiotic and seeded in 96-well plates at densities allowing them to reach confluence within four days. Two days after seeding, cells were transfected with GFP (100 ng/well; 2-3 rows; 16-24 wells or technical replicates) or one of five Hsa21 genes (100 ng/well; 1 row, 8 wells or technical replicates per unique cDNA) using Lipofectamine 2000 (Invitrogen, 11668030) according to manufacturer’s instructions. On day four, media was refreshed with DMEM containing 0.5% calf serum and 100 nM or 1 μM SAG. After forty-eight hours, cells were lysed and Fluc/Rluc activity was quantified using the Dual-Luciferase Reporter Assay System (Promega, E1910) and a 1450 MicroBeta Luminescence Counter (PerkinElmer). For the SmoA1-LIGHT screen, cells were seeded in 96-well plates at densities that would allow them to reach confluence within two days. One day after seeding, cells were transfected overnight with GFP or one of five Hsa21 genes and then switched to 0.5% calf serum media for twenty-four hours before quantification of Fluc/Rluc activity.

For both Shh-LIGHT2 and SmoA1-LIGHT screens, the Fluc/Rluc activity was normalized to the median value of the 96-well plate (intra-plate median centering). This process: 1) takes into account differences in the absolute intensity values between plates, 2) controls for unintended spatial gradients within plates, such as those that occur along the periphery, and 3) buffers against the presence of signaling outliers. Normalized values were then averaged for each Hsa21 cDNA or control gene. At minimum, all experiments were conducted with sets of 8 transfected wells. Technical replicates were averaged and Z-scores were calculated for each cDNA (*32*).

For validation studies of Shh-LIGHT2, cells were cultured to confluency in 96-well plates then treated with 1 uM SAG, 1 uM fluocinonide (Sigma SML0099), 1 uM fluticasone (Sigma, F9428), or 10 uM vitamin D3 (Sigma, C9756) in DMEM containing 0.5% calf serum.

### C3H10T1/2 differentiation

To quantify osteoblast differentiation following transfection of Hsa21 cDNAs, 5k C3H10T1/2 cells were seeded into each well of a 96-well plate. Twenty-four hours later, cells were transfected with 100 ng plasmid DNA per well using Lipofectamine 3000 according to manufacturer’s instructions (Invitrogen, L3000008). The position of each cDNA was randomized between experiments to minimize positional effects. Transfection efficiency was monitored in live cells via GFP expression. Twenty-four hours after transfection, cells were treated with plain media, 200 nM SAG, 2 uM cyclopamine (Calbiochem, 239806), or 200 nM SAG plus 2 uM cyclopamine. Four days after treatment, cells were washed with PBS and lysed with 50 ul passive lysis buffer (Promega, E194A) for 45 minutes. To quantify alkaline phosphatase activity, 200 ul alkaline phosphatase blue microwell substrate (Sigma, AB0100) was added to each well, and the plate was incubated in the dark for 30 minutes. Color development was measured using a SpectraMax 340 Microplate Reader (Molecular Devices) at 600 nm.

cDNAs were screened in two sets for a total of twenty independent replicates per chromosome 21 cDNA. Alkaline phosphatase activity was normalized to the median value of each plate. Cell viability was assessed 48, 72, and 96 hours after transfection using the MTT Cell Proliferation Assay kit (ATCC, 30-1010K) according to manufacturer’s instructions.

To stain cells for alkaline phosphatase activity, cells were fixed with 10% neutral buffered formalin (Sigma, HT501320) for one minute, permeabilized with 0.05% Tween-20 (Sigma, P9416) in PBS, and labeled with BCIP/NBT alkaline phosphatase substrate (Sigma, B5655). Cells were counterstained with nuclear fast red (Amresco, 1B1369) and dehydrated before mounting.

### RNA-seq

RNA-seq was performed essentially as previously described (*42*). Briefly, RNA from P6 cerebella was extracted and library preparation was conducted using the NEBNext® Poly(A) mRNA Magnetic Isolation Module (E7490) and NEBNext® Ultra™ II RNA Library Prep Kit for Illumina® (E7770). Library quality was assessed with an Agilent 2100 Bioanalyzer. Libraries were sequenced by the Johns Hopkins Single Cell & Transcriptomics Core (NovaSeq SP run, 50 bp paired-end reads) for an average of ∼54 million reads per sample.

Sequencing reads were mapped to the mouse genome mm39 modified by appending human chromosome 21, using the alignment tool STAR v.2.4.2a (*85*). The aligned reads were assembled with PsiCLASS v.1.0.2 (*86*) to create gene and transcript models. Unlike traditional transcript assemblers that process each sample separately, PsiCLASS simultaneously analyzes all samples in the experiment to produce a unified set of transcript annotations to use in the subsequent differential analyses. Transcripts were then assigned to known reference genes from the NCBI RefSeq databases (mouse release October 2020 and human release May 2021). Lastly, DESeq2 (*87*) was used to quantify the expression levels and determine the sets of differentially expressed genes. Additional visualizations, including plots of principal coordinate analysis (PCA) components, were visualized with custom R scripts. For comparison of human and mouse orthologs in the TcMAC21 model, trisomic counts were first length normalized using the formula len_norm_readcounts = 50 x readcounts/genelen. Human and mouse counts for each gene were then summed, and fold changes were reported as a ratio of TcMAC21 counts to euploid. Gene ontology and gene set enrichment analyses were performed using the R packages gprofiler2 v.0.2.1 (*88*), GSVA v.1.42.0 (*89*), GSEABase v.1.56.0, and clusterProfiler v.4.4.1 (*90*). Canonical pathways (reactome) gene set for gene set enrichment analysis was retrieved using the msigdbr R package v.7.4.1. Other R packages used to analyze and visualize RNA-seq data include tidyverse v.1.3.1 (*91*), cowplot v.1.1.1, ggbreak v.0.0.9 (*92*), ggrepel v.0.9.1, RColorBrewer v.1.1-2, gplots v.3.1.3, and enrichplot v.1.14.2 with scripts from DIY.transcriptomics (*93*).

### Lentiviral production

750k low-passage 293FT cells were seeded into each well of a 6-well plate coated with poly-L-ornithine (Sigma, P2533). One day after seeding, cells were transfected with 640 ng pMD2.G, 975 ng psPAX, and 1275 ng lentiviral target plasmid using Lipofectamine 2000 and PLUS reagent (Invitrogen, 11514015). Media was refreshed four hours after transfection. Supernatant was collected 48 and 72 hours post transfection, filtered with a 0.45 um filter (Millex-HV, SLHV013SL), and concentrated with Lenti-X Concentrator (Takara, 631231) according to manufacturer’s instructions. Physical titer was determined using the Lenti-X p24 Rapid Titer Kit (Takara, 632200) and granule cell precursors were transduced at an estimated MOI of ∼4.

### Granule cell precursor isolation

Cerebella from P6 pups were dissected into ice-cold Hanks’ Balanced Salt Solution (HBSS; Gibco, 14170112) with 0.6% glucose, digested with papain (Worthington Papain Dissociation System, LK003150), and triturated with a serum-coated pipette (*94*). Dissociated cells were isolated from membrane fragments on an albumin-ovomucoid inhibitor discontinuous density gradient. Granule cell precursors were further purified on a 35%/60% Percoll gradient (Sigma, E0414). Viable cells were counted with a Countess II Automated Cell Counter (ThermoFisher, A27978), and 100k cells were seeded into each well of a 96-well plate coated with poly-L-lysine (Sigma, P4832).

### Granule cell precursor EdU incorporation assay

Twenty-four hours after seeding, granule cell precursors were transduced with lentiviral particles. Infection was monitored via expression of GFP from the IRES-GFP construct. One day after transduction, media was refreshed with neurobasal media containing 6 nM SAG. Two days after transduction, cells were treated with 15 uM EdU for twenty-four hours. EdU incorporation was quantified using the Click-iT™ EdU Proliferation Assay for Microplates kit (Invitrogen, C10499) according to manufacturer’s instructions.

### Statistical analysis

Statistical analyses were performed using GraphPad Prism 9.1.2 or R version 4.1.3. For luciferase screens, z-scores were calculated by comparing the Fluc/Rluc ratio for each cDNA to the set of all screened cDNAs. For C3H10T1/2 alkaline phosphatase screen, non-parametric Kruskal-Wallis test was followed by Dunn’s post-hoc test comparing GFP control to all other cDNAs. All other assays were analyzed with two-tailed unpaired Student’s t-test, one-way ANOVA, or two-way ANOVA as noted. P-values were corrected for multiple comparisons by controlling the false discovery rate using the two-stage linear step-up procedure of Benjamini, Krieger and Yekutieli.

## Supplementary Materials

Fig. S1. Shh-LIGHT2 and SmoA1-LIGHT screens.

Fig. S2. C3H10T1/2 osteoblast differentiation and viability following transfection of chromosome 21 cDNAs.

Fig. S3. Expression of disomic genes in Ts65Dn and TcMAC21 cerebellum.

Fig. S4. Unsupervised clustering of differentially expressed genes in Ts65D and TcMAC21 samples.

Fig. S5. Gene ontology and gene set enrichment analyses of differentially expressed genes in Ts65Dn cerebellum.

Fig. S6. Differentially expressed genes in Ts65Dn cerebellum are implicated in human neurodevelopmental disorders, mitosis, and chromatin remodeling.

Fig. S7. Summary of effects of *HMGN1*, *MIS18A*, *B3GALT5*, and *ETS2* on SHH pathway activation.

Table S1. Cerebellar Volumes of Down Syndrome Models

Table S2. Manual Annotation of HSA21 Genes

Table S3. Plasmid Information

Table S4. Shh-LIGHT2 Screen

Table S5. SmoA1-LIGHT Screen

Table S6. Screen Comparisons

Table S7. Ts65Dn RNA-seq in P6 Cerebellum

Table S8. TcMAC21 RNA-seq in P6 Cerebellum

Table S9. Gene Set Enrichment Analysis in Ts65Dn Cerebellum

Table S10. Summary of Expression Data in Human and Mouse Cerebellum

## Acknowledgments

We would like to thank William Kaelin, Eric Campeau, Jesse Boehm, William Hahn, David Root, Aimin Liu, and Didier Trono for sharing plasmids and the Johns Hopkins Single Cell & Transcriptomics Core for providing sequencing services.

## Author contributions

Conceptualization: RHR, FXF, AJM

Formal analysis: AJM, LDF, YL, RHR

Funding acquisition: RHR, AJM

Investigation: AJM, FXF, YL, DKK

Project administration: RHR, AJM

Resources: DKK, AJM, YK, MO

Software: LDF

Supervision: RHR

Visualization: AJM, FXF

Writing – original draft: AJM

Writing – review & editing: RHR

## Data and materials availability

Plasmids are available from Addgene (https://www.addgene.org/Roger_Reeves/). Raw sequence data are deposited in the Gene Expression Omnibus, GEO accession number GSE202938. TcMAC21 mice are available from The Jackson Laboratory and require an agreement with RIKEN BRC and The National University Corporation Tottori University before shipping.

## Supplementary Figures

**Fig. S1.**
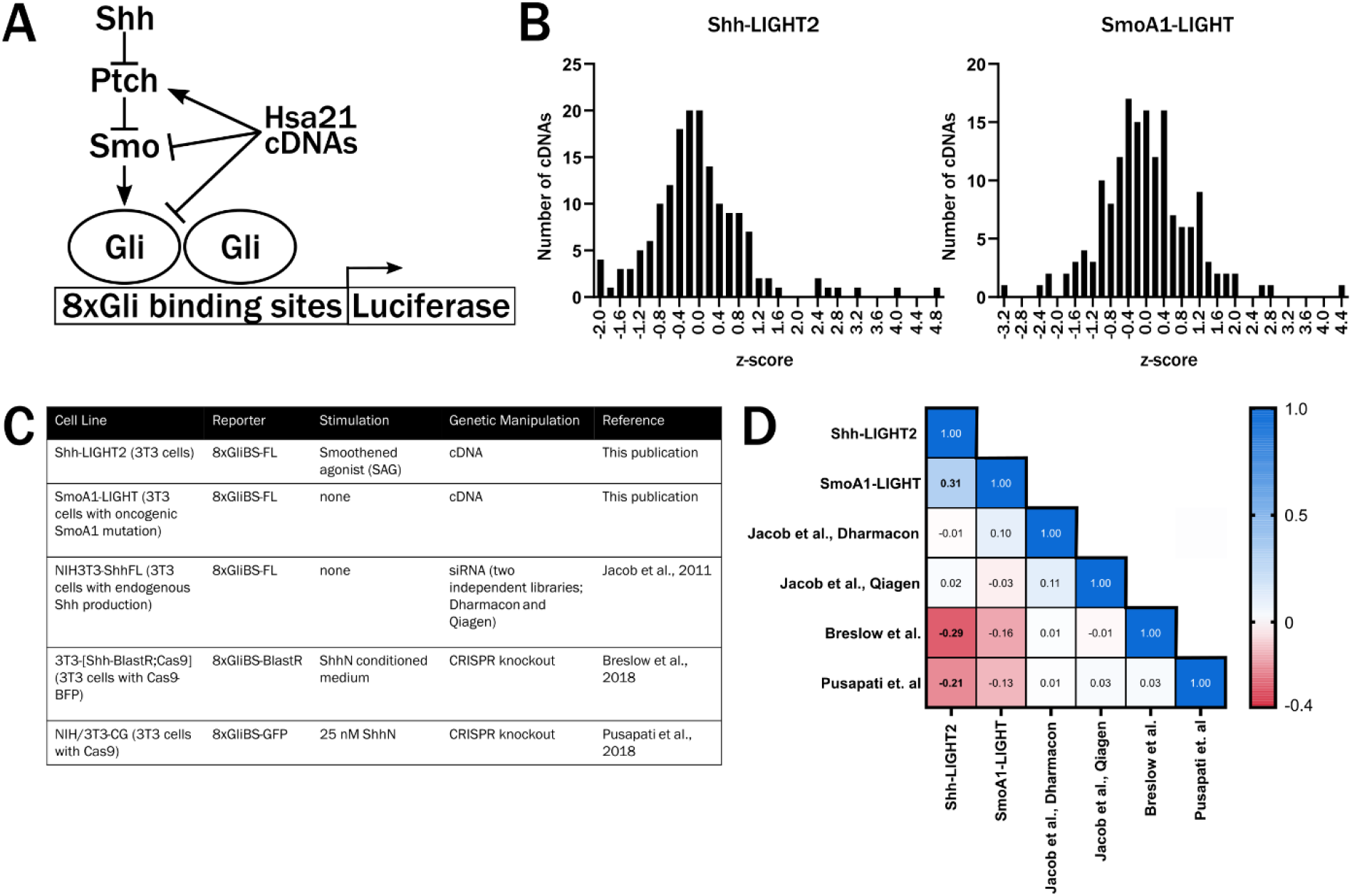
Shh-LIGHT2 and SmoA1-LIGHT screens. **(A)** Schematic of SHH signaling and 8xGliBS-FL reporter. Sonic hedgehog binds to Patched and relieves inhibition of Smoothened, which acts as a transducer to activate signaling via the Gli transcription factors. Binding of Gli to the 8xGliBS-FL promotes transcription of luciferase. Overexpression of chromosome 21 genes may activate or inhibit SHH signaling at any level of the signaling pathway. **(B)** Distribution of z-scores in Shh-LIGHT2 and SmoA1-LIGHT cDNA overexpression screens. **(C)** Summary of previously reported siRNA knockdown and CRISPR knockout screens using the 8xGliBS reporter. **(D)** Correlation matrix showing Pearson correlation coefficient r between pairs of screens for the 115 chromosome 21 genes and mouse orthologs with data across all screens. Bolded correlation coefficients have P < 0.05. The Shh-LIGHT2 and SmoA1-LIGHT cDNA screens are positively correlated, whereas Shh-LIGHT2 shows a negative correlation with two CRISPR knockout screens.

**Fig. S2.**
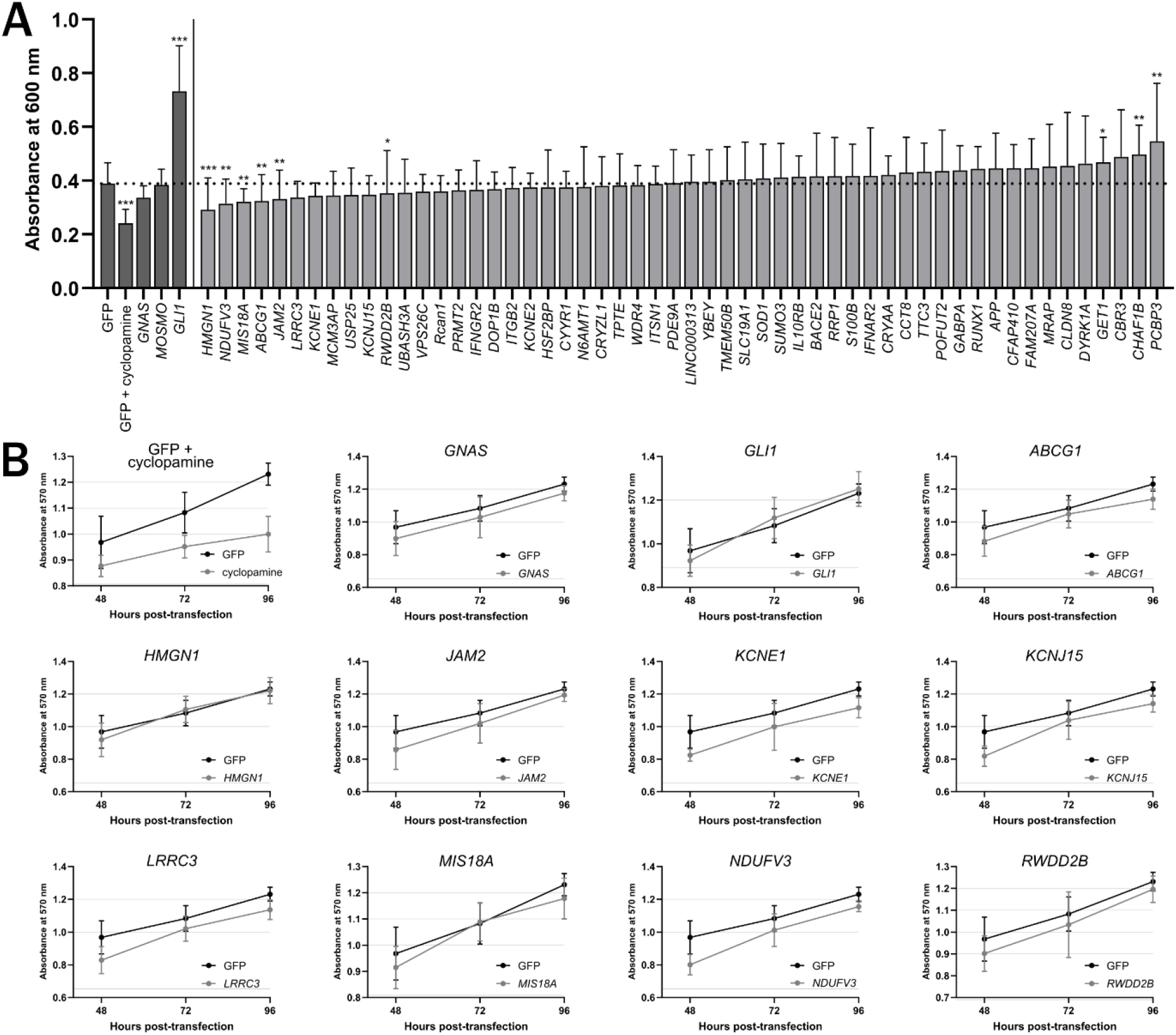
C3H10T1/2 osteoblast differentiation and viability following transfection of chromosome 21 cDNAs. **(A)** Quantification of alkaline phosphatase activity following transfection of chromosome 21 cDNAs and stimulation with SAG (n=20). All conditions were compared to GFP control. ***P<0.001, **P<0.01, *P<0.05 (Kruskal-Wallis test followed by Dunn’s post-hoc test). **(B)** Quantification of viability of untreated C3H10T1/2 cells 48, 72, and 96 hours post-transfection (n=7). In cells treated with SAG, only cyclopamine treatment affected viability (data not shown).

**Fig. S3.**
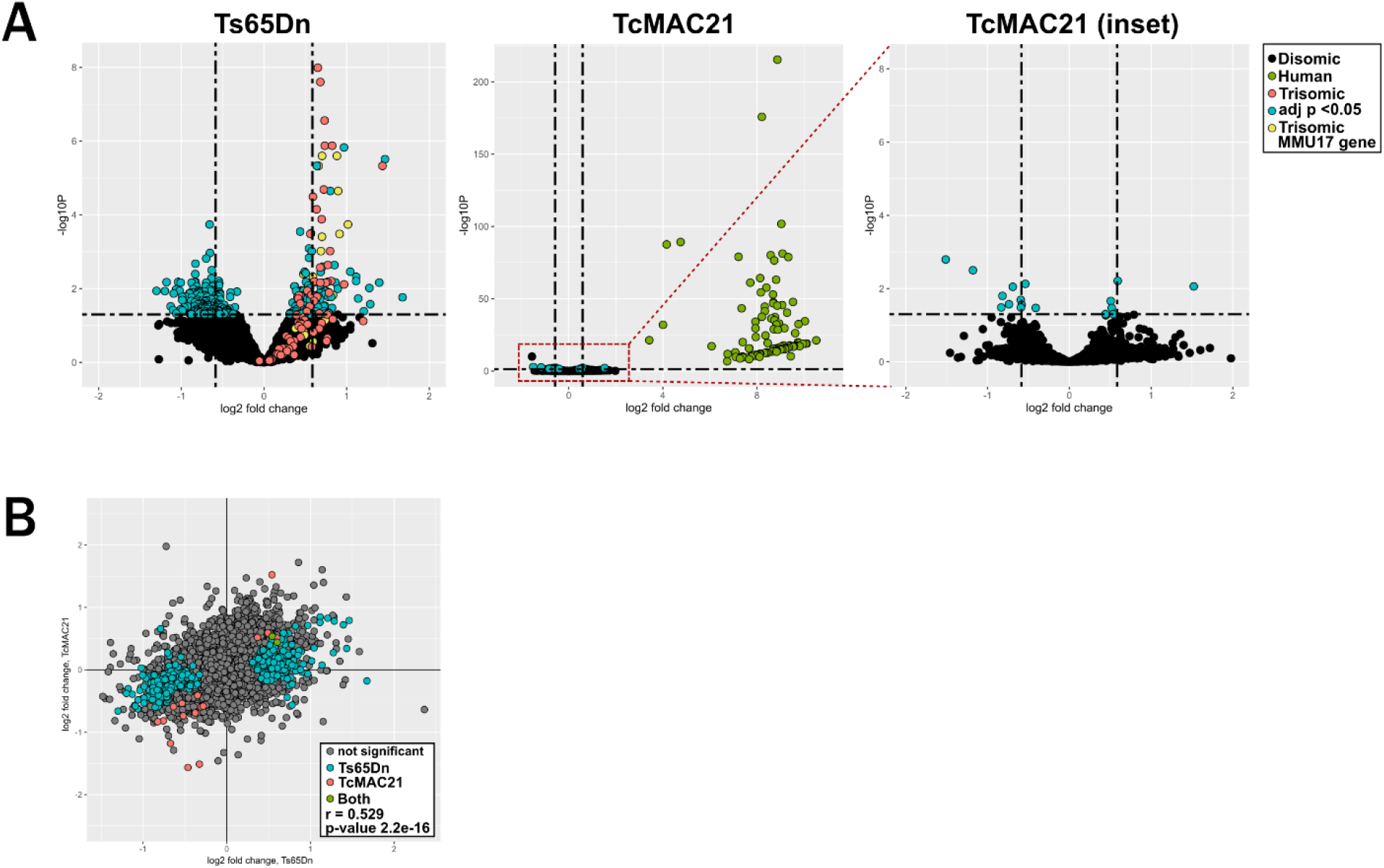
Expression of disomic genes in Ts65Dn and TcMAC21 cerebellum. **(A)** Volcano plots showing log2 fold change and −log10(P value) in Ts65Dn and TcMAC21 samples. Teal points represent disomic transcripts with adjusted P <0.05, salmon points represent chromosome 21 orthologs that are trisomic in Ts65Dn, yellow points represent non-chromosome 21 orthologs (MMU17) transcripts that are trisomic in Ts65Dn, and green points represent human transcripts in TcMAC21 samples. **(B)** Scatterplot showing log2 fold change of disomic transcripts in Ts65Dn and TcMAC21 samples. Teal points are significantly differentially expressed in Ts65Dn samples, salmon points are significantly differentially expressed in TcMAC21 samples, and green points are differentially expressed in both Ts65Dn and TcMAC21 samples. Pearson correlation coefficient r=0.529 and P=2.2e-16.

**Fig. S4.**
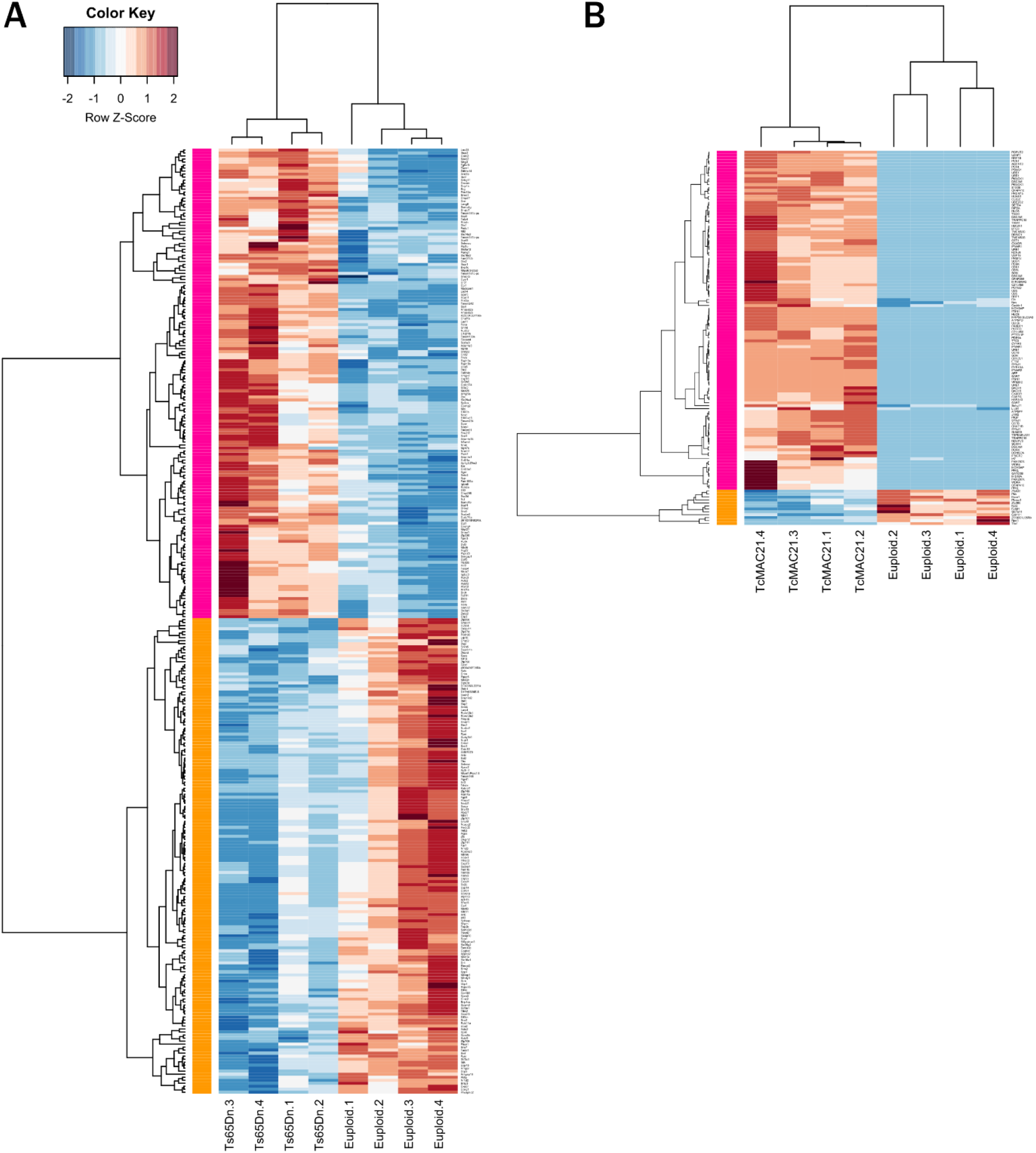
Unsupervised clustering of differentially expressed genes in Ts65D and TcMAC21 samples. **(A)** Unsupervised clustering of 314 differentially expressed transcripts (rows) in 4 Ts65Dn cerebella and 4 euploid littermates (columns). The orange module represents genes downregulated in Ts65Dn relative to control and the pink module represents genes upregulated in Ts65Dn. **(B)** Unsupervised clustering of 127 differentially expressed transcripts in 4 TcMAC21 cerebella and 4 euploid littermates. 109/127 differentially expressed transcripts derive from the HSA21q-MAC hybrid chromosome.

**Fig. S5.**
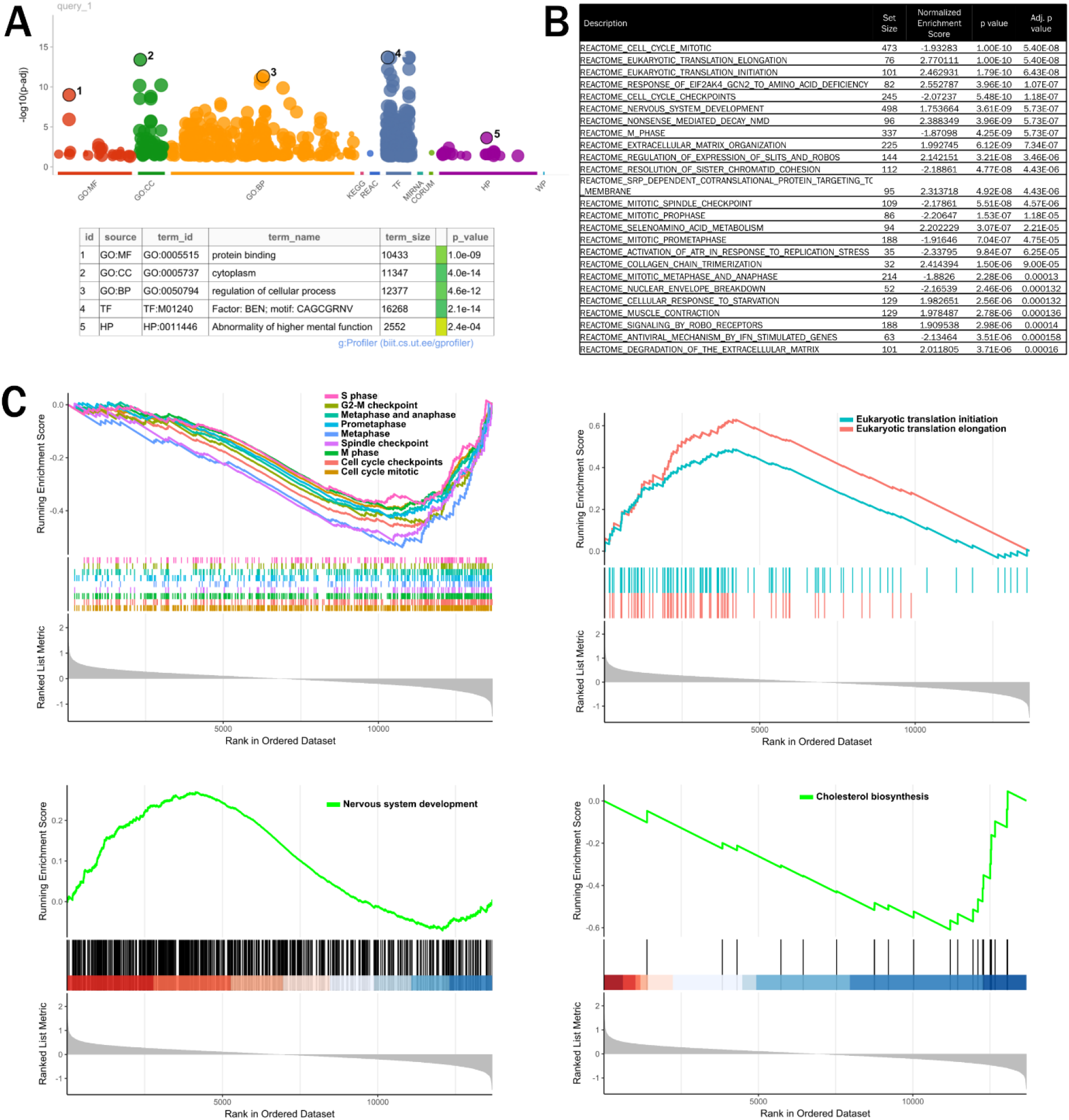
Gene ontology and gene set enrichment analyses of differentially expressed genes in Ts65Dn cerebellum. **(A)** Manhattan plot showing top gene ontology terms identified in Ts65Dn cerebellum. Differentially expressed genes contributing to “abnormality of higher mental function” are listed in figure S6. **(B)** Top 25 pathways identified by gene set enrichment analysis in Ts65Dn samples. **(C)** Gene set enrichment analysis for pathways significantly enriched in Ts65Dn samples (translation and nervous system development) and pathways enriched in control samples (mitotic/cell cycle and cholesterol biosynthesis).

**Fig. S6.**
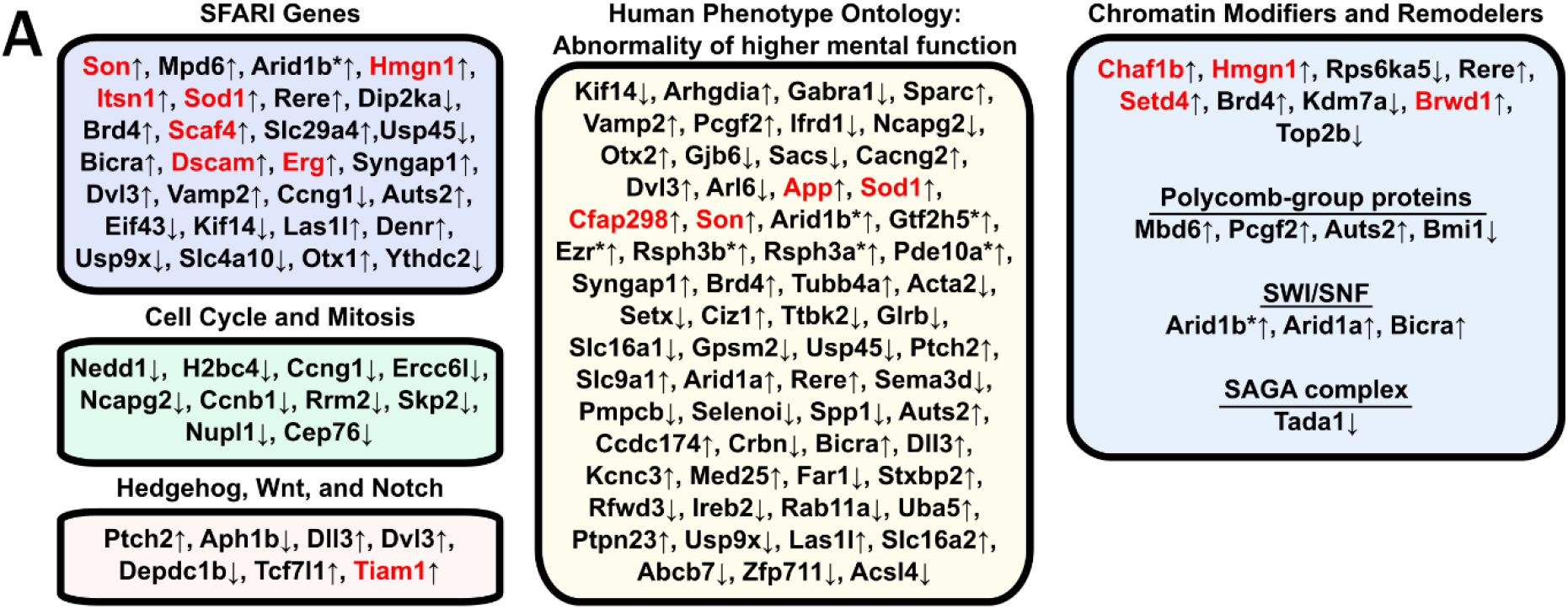
Differentially expressed genes in Ts65Dn cerebellum are implicated in human neurodevelopmental disorders, mitosis, and chromatin remodeling. **(A)** Subset of differentially expressed genes (q < 0.05) identified in Ts65Dn grouped by cellular and disease processes. Up arrows signify genes that are upregulated in Ts65Dn samples, asterisks signify genes that are trisomic in Ts65Dn mice but are not orthologs of chromosome 21 genes, and red signifies trisomic genes that are orthologs of chromosome 21 genes. Differentially expressed genes include 28 in the SFARI Gene database of autism susceptibility loci and others related to cell cycle/mitosis, key developmental pathways including SHH, and chromatin modifiers and remodelers.

**Fig. S7.**
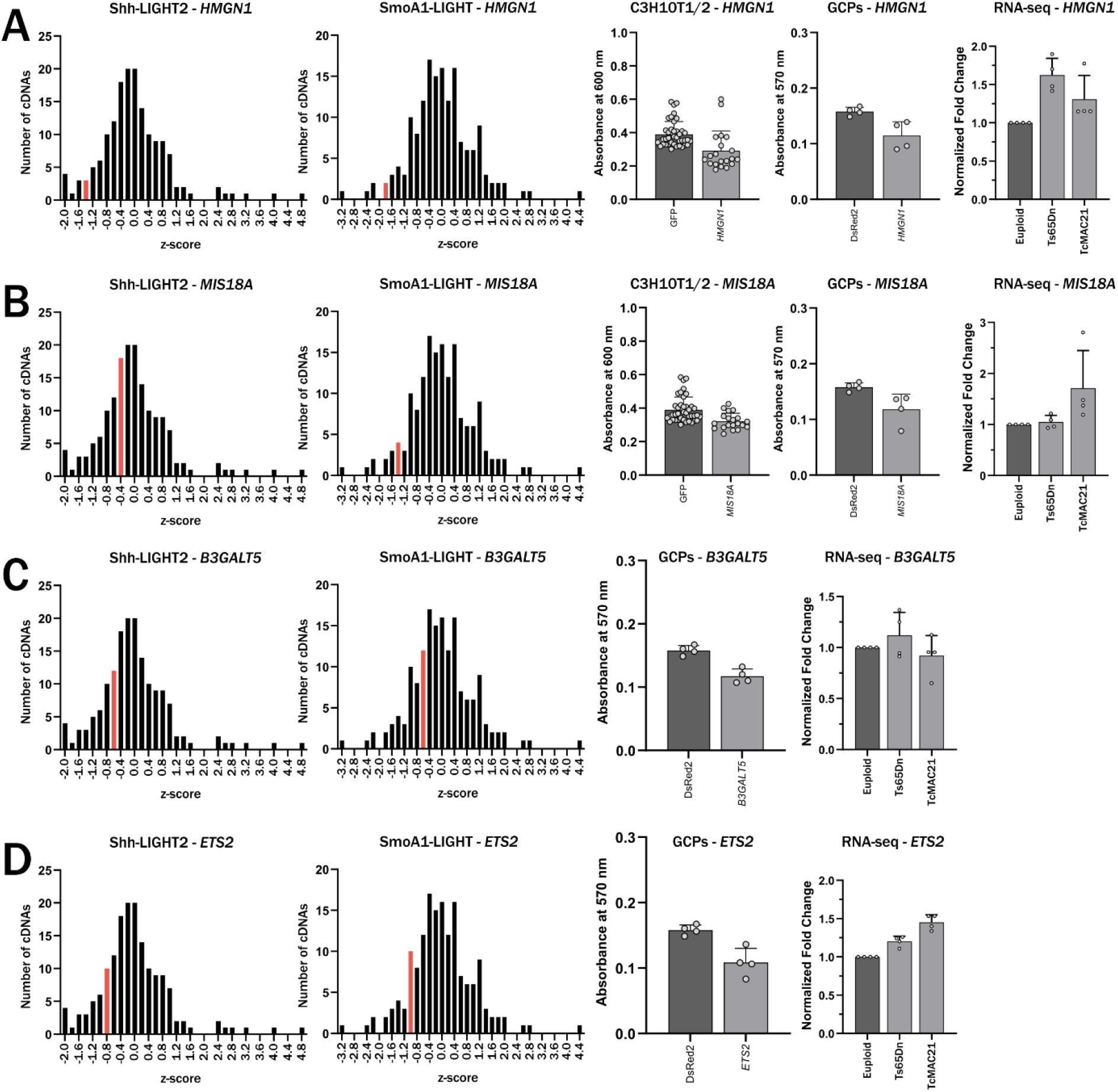
Summary of effects of *HMGN1*, *MIS18A*, *B3GALT5*, and *ETS2* on SHH pathway activation. **(A)** Effect of *HMGN1* overexpression on luciferase activity in Shh-LIGHT2 and SmoA1-LIGHT cells, osteoblast differentiation in C3H10T1/2 cells, and proliferation in primary granule cell precursors. Bin containing *HMGN1* for Shh-LIGHT2 and SmoA1-LIGHT assays is highlighted in red. Overexpression of *HMGN1* or its mouse ortholog in Ts65Dn and TcMAC21 cerebellum graphed relative to euploid controls (n=4). Same for **(B)** *MIS18A*, **(C)** *B3GALT5*, and **(D)** *ETS2*.

